# Characterizing receptor flexibility to predict mutations that lead to human adaptation of influenza hemagglutinin

**DOI:** 10.1101/2020.07.15.204982

**Authors:** Huafeng Xu, Timothy Palpant, Cody Weinberger, David E. Shaw

## Abstract

A key step in the emergence of human pandemic influenza strains has been a switch in the binding preference of the viral glycoprotein hemagglutinin (HA) from avian to human sialic acid receptors (SAs). The conformation of the bound SA varies substantially with HA sequence, and crystallographic evidence suggests that the bound SA is flexible, so it is difficult to predict from crystal structures which mutations are responsible for the change in HA binding preference. We performed molecular dynamics (MD) simulations of SA analogs binding to various HAs, and observed a dynamic equilibrium among structurally diverse receptor conformations, including novel conformations that have not been experimentally observed. Using one such novel conformation, we predicted—and subsequently confirmed with microscale thermophoresis experiments—a set of mutations that substantially increased an HA’s affinity for a human SA analog. This prediction could not have been inferred from existing crystal structures, suggesting that MD-generated HA-SA conformational ensembles could help researchers predict human-adaptive mutations, aiding in the surveillance of emerging pandemic threats.

---

For a strain of influenza to become pandemic, a non-human—often avian—strain must acquire mutations that enable it to infect and transmit between humans.^1,2,3^ The infection of a host cell by an influenza virus begins when the glycoprotein hemagglutinin (HA) on the viral surface binds to HA-specific receptors (i.e., sialic acid–containing glycoproteins and glycolipids) on the host cell.^4^ Human transmissibility requires both that the viral HAs have sufficiently strong affinity for the human sialic acid receptors (SAs) on the target cells to establish adequate adhesion,^5,6^ and that they have sufficiently weak affinity for respiratory tract mucins to avoid sequestration and reach the target cells.^7^ Human-transmissible strains are associated with mutations in HA that switch its binding specificity from avian to human receptors.^8–13^ Such mutations typically increase the binding affinity to receptors whose terminal sialic acid is linked to the penultimate sugar by an α2,6 glycosidic bond (prevalent on epithelial cells in the human trachea, the site of human infection) and decrease binding affinity to receptors whose terminal sialic acid uses an α2,3 linkage (prevalent in respiratory tract mucins and in avian enteric tracts, the latter being the primary site of avian infection).^14,15^

Various distinct sets of mutations have caused HAs of different subtypes, which have substantially different amino acid sequences^16^ (Fig. 1), to switch binding preference from avian to human SAs in past pandemics.^17^ Although X-ray crystallography, biochemical studies, and computational studies^18–20^ have shed some light on the structural and thermodynamic basis of these switching mutations,^9,21–28^ it has been a challenge in studies of past affinity-switching strains to identify the key sequence differences between subtypes that allowed a given set of mutations to confer human-SA binding preference to HA of one subtype but not another, and it remains difficult to predict which sets of mutations would enable an HA of a given sequence to switch binding preference in the future.^29^

**Figure 1.**
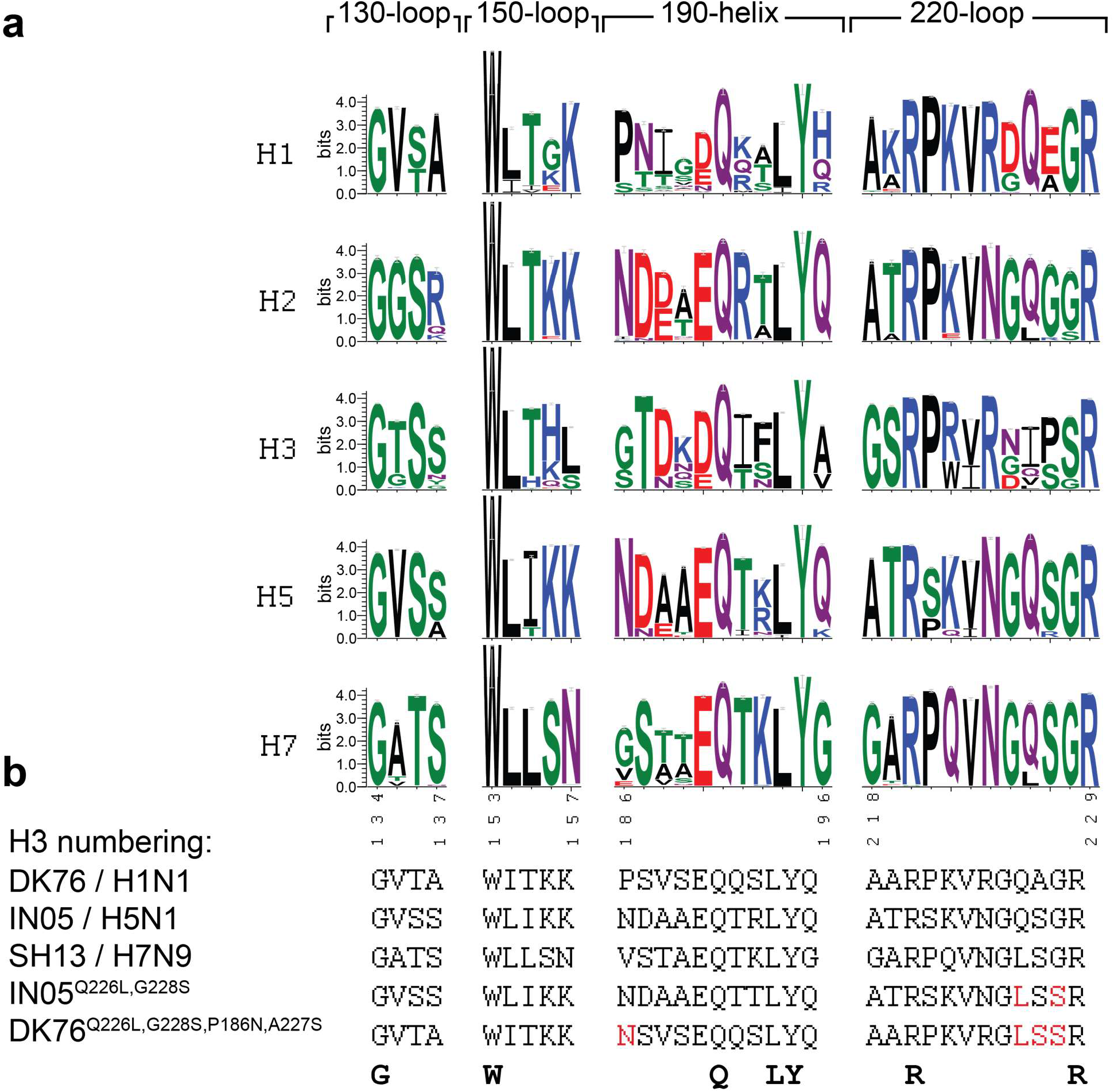
The multiple-sequence alignment (MSA) is shown for the 130-loop, the 150-loop, the 190-helix, and the 220-loop, which form the binding pocket, in HAs of different subtypes. (**a**) Sequence logo representation of HAs of H1, H2, H3, H5, and H7 subtypes. The height of the letter indicates the frequency that the corresponding amino acid is observed at that sequence position. There is substantial sequence variation in the binding pocket within a subtype and even more difference across different subtypes. (**b**) The sequences of the HAs of the DK76, IN05, and SH13 strains, and some of the variants studied in this work. Highlighted in red are the mutations that enabled the HA to switch binding preference from avian receptors to human receptors. In the H1N1 pandemic strain, the pair of mutations E190D,G225D in HA led to the switch in binding preference,^34^ whereas in the H2N2 and H3N2 pandemic strains, the pair of mutations Q226L,G228S caused the switch. The latter pair of mutations can also confer preferential binding to human receptors when introduced into HA of the H5N1 IN05 strain.^32,33^ No HA of H2 or H3 subtypes, however, has been reported to switch binding preference with the E190D,G225D mutations, and this pair of mutations has been shown not to switch the binding preference when introduced into an HA of H5 subtype.^8^ Conversely, no HA of the H1 subtype has been reported to switch its binding preference by the Q226L,G228S mutations; an additional pair of mutations A227S,P186N, identified in this work, were required for DK76 HA to do so. The full sequences of DK76, IN05, SH13, and their mutants studied in this work are provided in the SI.

This difficulty arises in part because of the conformational heterogeneity of SAs when bound to different HA variants: Although the terminal N-acetylneuraminic acid (Neu5Ac) adopts the same conformation in all human and avian HA-SA complexes (Figs. S1a and S1b), the linked penultimate galactose (Gal) and the third sugar—often N-acetylglucosamine (GlcNAc)—of SAs can assume many different conformations when bound to different HA variants (Fig. S2). Switching mutations in past pandemic strains have primarily affected HA interactions with Gal and GlcNAc (Figs. S1c and S1d), and it is thus likely that predicting affinity-switching mutations in HA would require knowing the range of possible binding conformations of the SA’s terminal three sugars.

Furthermore, the GlcNAc and Gal moieties of human SAs are only partially resolved in many crystal structures of HA-SA complexes, hinting that human receptors might remain flexible and adopt a wide range of conformations while bound to HA. In existing crystal structures, human SAs adopt a more diverse range of binding conformations across complexes with different HAs than avian SAs do (Fig. S2), suggesting that human receptors may be especially flexible. It would be difficult, however, to structurally characterize flexible SA conformations in a given HA-SA complex using X-ray crystallography.

Here we report the results of long-timescale molecular dynamics (MD) simulations^30,31^ of binding between a number of HA variants and analogs of SAs. In our simulations, we observed that human SAs in complex with HA remained in a dynamic equilibrium among a diverse ensemble of conformations, adopting both all of the crystallographically determined binding conformations and a number of novel binding conformations. Based on one of these novel binding conformations, we predicted the effects of a number of previously unexamined HA mutations on HA-SA binding affinity, including a set of mutations that we predicted would substantially increase the binding affinity of an avian HA for a human receptor analog. These predictions, which could not have been made using existing crystal structures, were then tested using microscale thermophoresis (MST) experiments. The results of these experiments were consistent with our predictions. Our results thus suggest that ensembles of binding conformations generated by MD might be used by researchers to help predict potential human-adaptive mutations in avian hemagglutinin, which could potentially assist in the monitoring of future pandemic threats.

We simulated the binding of α2,3-linked sialyl lactosamine (3-SLN, Fig. S1e), a trisaccharide analog of avian receptors, and of α2,6-linked sialyl lactosamine (6-SLN, Fig. S1f), a trisaccharide analog of human receptors, to a diverse set of variants (Table S1) of HAs from three different subtypes: A/duck/Alberta/35/1976 (DK76) of the H1 subtype, A/Indonesia/5/2005 (IN05) of the H5 subtype, and A/Shanghai/2/2013 (SH13) of the H7 subtype. We also simulated the binding of another human receptor analog, the pentasaccharide 506-SLN (also known as LSTc), to a subset of the HA variants. (We denote an HA variant by its strain of origin and, in the superscript, its amino acid mutations from the wild type; a mutation is abbreviated by the one-letter code of the amino acid in the wild type, followed by the HA residue number according to the numbering in the H3 subtype, followed by the one-letter code of the amino acid in the mutant). The receptor analogs 3-SLN, 6-SLN, and 506-SLN represent the receptors’ terminal sugars that interact directly with HA, and they have been commonly used in structural and biochemical studies of receptor-HA binding.

In our simulations, 3-SLN and 6-SLN—and 506-SLN for some HA variants—spontaneously bound to and unbound from HA (Figs. 2a, 2b, and S3; Movie S1); when 3-SLN, 6-SLN, and 506-SLN were bound to an HA molecule, they recapitulated the crystallographic binding conformation of Neu5Ac and the interactions conserved across all HA-SA complexes. In every one of the simulated HA-SA complexes, the parts of the SAs other than Neu5Ac remained flexible and adopted a wide range of conformations, including novel conformations previously unseen in crystal structures of any HA-SA complexes.

**Figure 2.**
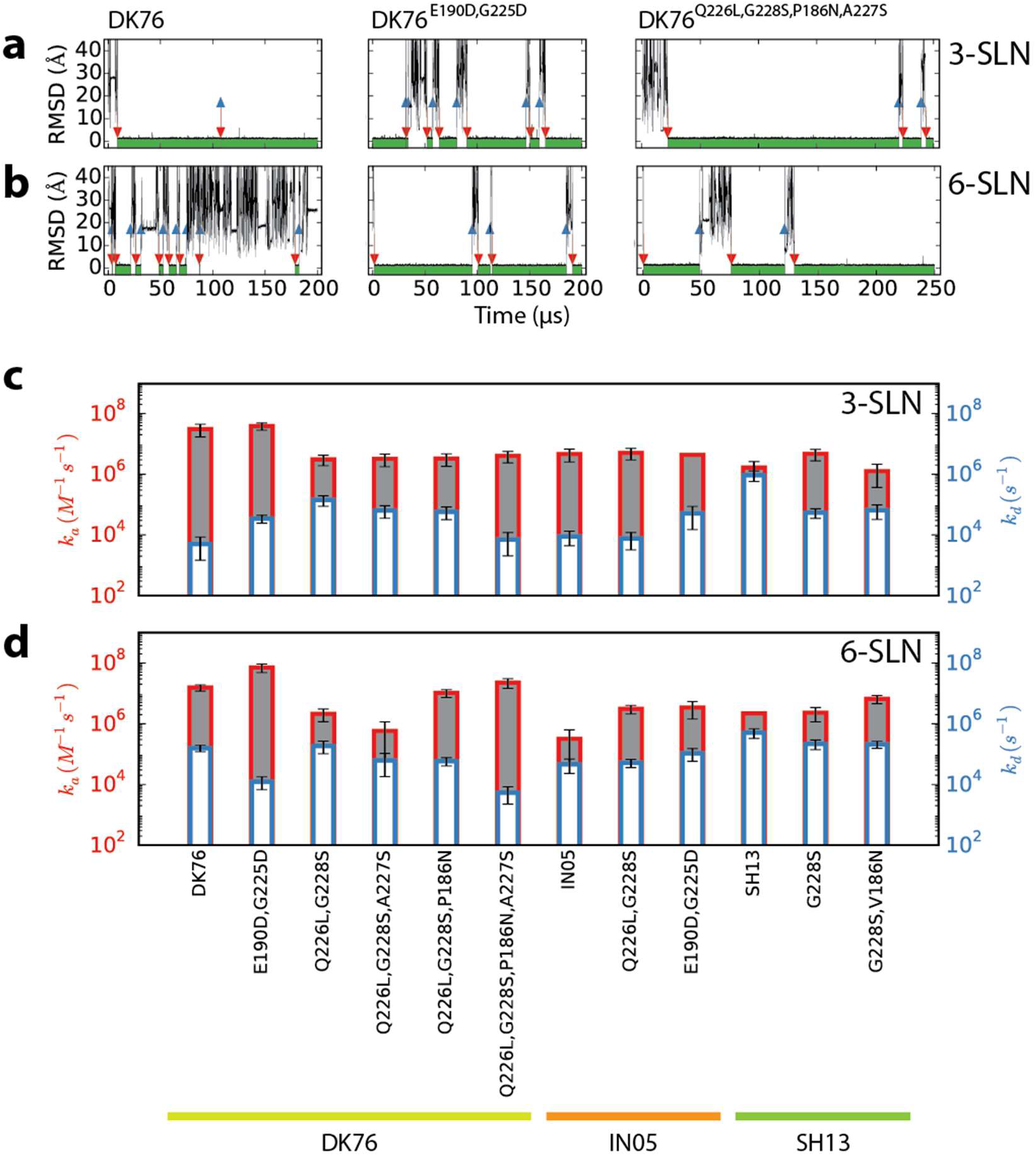
Receptor analogs reversibly bound to and unbound from HA in MD simulations, and the kinetic rates can be estimated from the binding and unbinding times. (**a, b**) Representative time traces of the root-mean-square deviation (RMSD) of Neu5Ac from its bound pose in crystal structures, in the simulations of (**a**) 3-SLN and (**b**) 6-SLN binding to the RBD of the DK76 HA and two of its mutants. The times of binding (i.e., when the RMSD fell below 1.5 Å) and unbinding (i.e., when the RMSD rose above 8 Å) are indicated by red and blue arrows, respectively, and the time intervals when the receptor analog was bound are indicated by the green shade under the curve. (**c, d**) The kinetic rates, *k*_*a*_ and *K*_*D*_, for (**c**) 3-SLN and (**c**) 6-SLN binding to different HA variants, estimated from the MD simulations. Since the rates are plotted on a logarithmic scale, the difference between *k*_*a*_ and *K*_*D*_, indicated by the height of the gray area, gives the equilibrium association constant. The rates and the statistical errors are estimated with the assumption that the binding and unbinding events followed Poisson statistics. The RMSD time traces of all of the MD simulations used to estimate the *k*_*a*_ and *K*_*D*_ values are shown in Fig. S3. Absence of the error bar on *k*_*a*_ indicates that the value is an upper bound, reflecting the fact that no spontaneous binding event occurred in the corresponding simulations, and that based on the lengths of the simulations we can thus estimate that there is only a P = 0.02 probability that *k*_*a*_ is greater than the given value (see SI for details).

To give confidence that the glycan force field parameters^43^ used in this work are sufficient to yield accurate conformational ensembles of SAs, we simulated the receptor analogs 6-SLN, 506-SLN, and 503-SLN (also known as LSTa), which have been previously characterized using nuclear magnetic resonance (NMR),^45,46^ in aqueous solution, and we compared the nuclear Overhauser effect (NOE) and *J*_3_-coupling parameters estimated from these simulations to the NMR measurements. The good agreement between the computational estimates and the experimental values (Tables S4–S6)—in addition to the fact that our simulations reproduced all known SA binding conformations observed in crystallography—suggests that the receptor conformations in our simulations are likely realistic.

From the binding and unbinding events in the long-timescale simulations, the kinetic rates of association (*k*_*a*_) and dissociation (*k*_*d*_), and the equilibrium dissociation constant (*K*_*D*_), can be estimated (Figs. 2c and 2d; Table 1; see SI for the estimation method). Although such predicted *K*_*D*_ values have large statistical uncertainties and may differ in absolute terms from experimental values (as discussed in more detail below), they are consistent with the previously observed mutational patterns of human adaptation: IN05 of the H5 subtype gains affinity to human receptors by the mutations Q226L,G228S,^32,33^ whereas DK76 of H1 subtype does not; the latter switches its binding preference from avian to human receptors by the mutations E190D,G225D.^34^ These results also suggest that our simulations are able to capture the changes in HA-SA interactions that result from HA mutations.

**Table 1.** The equilibrium dissociation constants (*K*_*D*_) of different receptor analogs binding to the IN05 and DK76 HA variants are shown. Only the RBD of HA was included in the simulations, whereas the full HA trimer was used in the MST assays. The *K*_*D*_ results from MST are thus apparent dissociation constants of the receptor analog binding to the HA trimer, whereas those from MD are dissociation constants of binding to the monomeric RBD. “N.B.” indicates no detectable binding in MST; blank cells indicate that the corresponding MD simulation or MST experiment was not performed. For the MD results, the statistical uncertainties are reported in the superscript and subscript, which correspond to the upper and lower bounds, respectively, of the 68.3% confidence interval of the estimates (i.e., at one standard error in the estimate of ln(*K*_*D*,MD_). The *K*_*D*,MST_ value for IN05^Q226L,G228S,S227A^ (with superscript “a”) was estimated from fluorescence quenching, as no binding was detectable from thermophoresis.

| hemagglutinin | receptor | $K_{D,MD}$ ($10^{-3}M$) | $K_{D,MST}$ ($10^{-3}M$) |
| --- | --- | --- | --- |
| IN05 | 3-SLN | $2^{5.4}_{0.74}$ | N.B. |
| | 6-SLN | $200^{544}_{74}$ | N.B. |
| | 506-SLN | | $0.35 \pm 0.02$ |
| IN05<sup>Q226L,G228S</sup> | 3-SLN | $2^{3.3}_{1.2}$ | N.B. |
| | 6-SLN | $20^{33}_{12}$ | $0.055 \pm 0.004$ |
| | 506-SLN | | $0.21 \pm 0.01$ |
| IN05<sup>Q226L,G228S,S227A</sup> | 6-SLN | $182^{344}_{96}$ | $1.6 \pm 0.1^a$ |
| DK76 | 3-SLN | $0.1^{0.3}_{0.04}$ | $0.64 \pm 0.06$ |
| | 6-SLN | $10^{16.5}_{6.1}$ | N.B. |
| | 506-SLN | $10^{10966}_{0.01}$ | $0.088 \pm 0.009$ |
| DK76<sup>E190D,G225D</sup> | 3-SLN | $0.9^{1.6}_{0.5}$ | N.B. |
| | 6-SLN | $0.1^{0.3}_{0.04}$ | $0.20 \pm 0.02$ |
| | 506-SLN | | $0.08 \pm 0.01$ |
| DK76<sup>Q226L,G228S</sup> | 3-SLN | $50^{91}_{27}$ | N.B. |
| | 6-SLN | $90^{219}_{37}$ | N.B. |
| | 506-SLN | $40^{85}_{19}$ | $0.11 \pm 0.01$ |
| DK76<sup>Q226L,G228S,P186N,A227S</sup> | 3-SLN | $2^{5.4}_{0.74}$ | $0.04 \pm 0.03$ |
| | 6-SLN | $0.2^{0.5}_{0.07}$ | $0.06 \pm 0.02$ |
| | 506-SLN | $30^{81.5}_{11}$ | $0.23 \pm 0.05$ |
| DK76<sup>Q226L,G228S,P186N</sup> | 3-SLN | $20^{54}_{7.4}$ | |
| | 6-SLN | $6^{9.9}_{3.6}$ | N.B. |
| DK76<sup>Q226L,G228S,A227S</sup> | 3-SLN | $10^{27}_{3.7}$ | |
| | 6-SLN | $100^{739}_{13.5}$ | N.B. |

When the human receptor analogs (6-SLN and 506-SLN) were bound to any of the simulated HA variants, the Gal and GlcNAc moieties assumed a diverse set of conformations. The receptors rapidly interconverted among structurally dissimilar binding conformations (Fig. 3a), such as the cis and trans configurations around the glycosidic linkage between Neu5Ac and Gal^9,22^ (Table S2). Each human receptor analog bound to any individual HA variant visited the majority of all the binding conformations that have been observed in the crystal structures of a variety of receptor analogs bound to diverse HA variants of different subtypes (Fig. S4). The binding conformations of the receptors in our simulations can be divided into distinct clusters (Figs. 3b, S5, S6, and S7). The occupancy in each cluster varied by HA variant (Figs. 3c and S7b) as well as between 6-SLN and 506-SLN. The dominant binding conformation in the simulations was not always the crystallographic conformation of the corresponding HA variant (Fig. S5). In addition to the conformations previously observed in crystal structures, both human receptor analogs also bound to HA variants in novel conformations (Figs. 3b and S7a). Human receptors displayed less conformational flexibility when bound to human-adapted HA mutants than to the corresponding wild-type avian HAs; a given human receptor bound to a human-adapted mutant predominantly occupied a single conformation, one which in some cases was only rarely visited in its complex with the wild-type avian HA.

**Figure 3.**
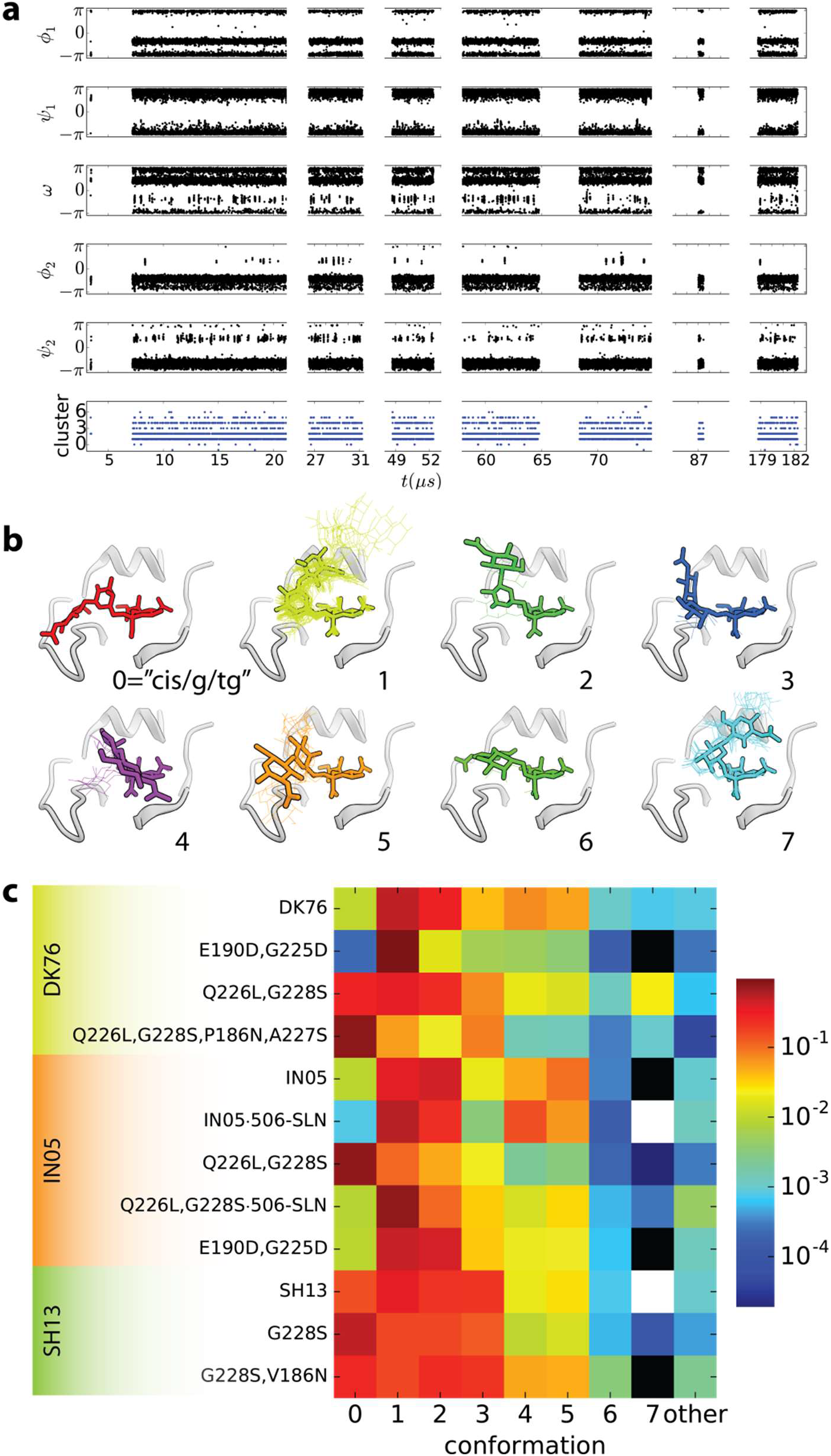
Receptor analogs adopted a diverse and dynamic ensemble of conformations when 25 bound to HA. To characterize these conformations, we projected each conformation onto five torsional angles: around the three bonds connecting Gal to Neu5Ac and around the two bonds connecting GlcNac to Gal (Figs. S1c and S1d). (**a**) The time trace of the five torsional angles (see Fig. S1f) and the conformational-cluster assignment in the simulation of 6-SLN in complex with DK76, showing the transitions between different conformations. Discontinuity in time reflects the intervals when 6-SLN unbound. (**b**) The binding conformations of the receptors in our simulations can be divided into 8 clusters based on the three torsional angles φ_1_, Ψ_1_, and ω, which determine the position of the Gal ring. The receptor conformation corresponding to the center of each cluster is shown here in thick sticks. Crystal structures, shown in thin lines, are assigned to the cluster with the smallest RMSD between its center and the crystal structure, provided that the RMSD is also smaller than 1.5 Å. The conformations in cluster 0 have not yet been observed in crystal structures. (**c**) The occupancy of different clusters of conformations by human receptor analogs bound to different HA variants. The occupancy varies among HA variants and between 6-SLN and 506-SLN.

The above results may partly explain the difficulty in predicting human-adapting mutations based on crystal structures. They suggest that, unlike other examples of specific biomolecular binding (which are normally associated with a unique, well-defined binding conformation), a given human receptor binds to each HA variant in a number of distinct binding conformations. Crystal structures of a receptor in complex with different HA variants may thus give an indication of the range of the receptor’s possible binding conformations in a fluid ensemble, as has been suggested for other biomolecular complexes.^35^

Next, we explore whether the heterogeneous binding conformations sampled in our MD simulations allow us to predict HA mutations that might affect binding to human receptors but cannot be inferred from crystal structures alone, hypothesizing that new favorable interactions between HA and SA might form in some of the predicted novel conformations. We use long-timescale MD simulations to estimate, and MST experiments^36^ to measure, the effects of the predicted mutations on the binding affinities between the receptor analogs and HA variants. (The sequences of the HA variants used in the MST experiments are listed in Table S3).

In this work, we examine the conformation of cluster 0, which we term the cis/g/tg conformation because its first torsional angles are *ϕ*_1_ ≈ −π / 3 (cis), *ϕ*_1_ ≈ −π / 2 (gauche), and ω ≈ π (trans-gauche). In this conformation, which to our knowledge has not previously been observed experimentally or in simulation, the receptor exits the binding pocket over the 220 loop (Fig. 4a). The glutamate at position 190 (E190) forms a hydrogen bond with the O_2_ hydroxyl of galactose, and in the IN05 and SH13 HA variants, the serine at position 227 (S227) forms a hydrogen bond with the terminal hydroxyl of GlcNAc in 6-SLN. These two interactions, which do not appear in any previously published crystal structure (Fig. 4b), may contribute to the increased affinity that the mutant IN05^Q226L,G228S^, from the mammal-transmissible H5 strain, has for 6-SLN compared to wild-type IN05 (Figs. 2c and 2d; Table 1).^9^ We thus predicted that mutating Ser 227 in IN05^Q226L,G228S^—a position where no mutation has been previously reported to affect receptor binding in any HA—to Ala would weaken binding to 6-SLN. Introducing the mutation S227A in IN05^Q226L,G228S^ indeed reduces its binding affinity for 6-SLN in our MD simulations (Table 1).

**Figure 4.**
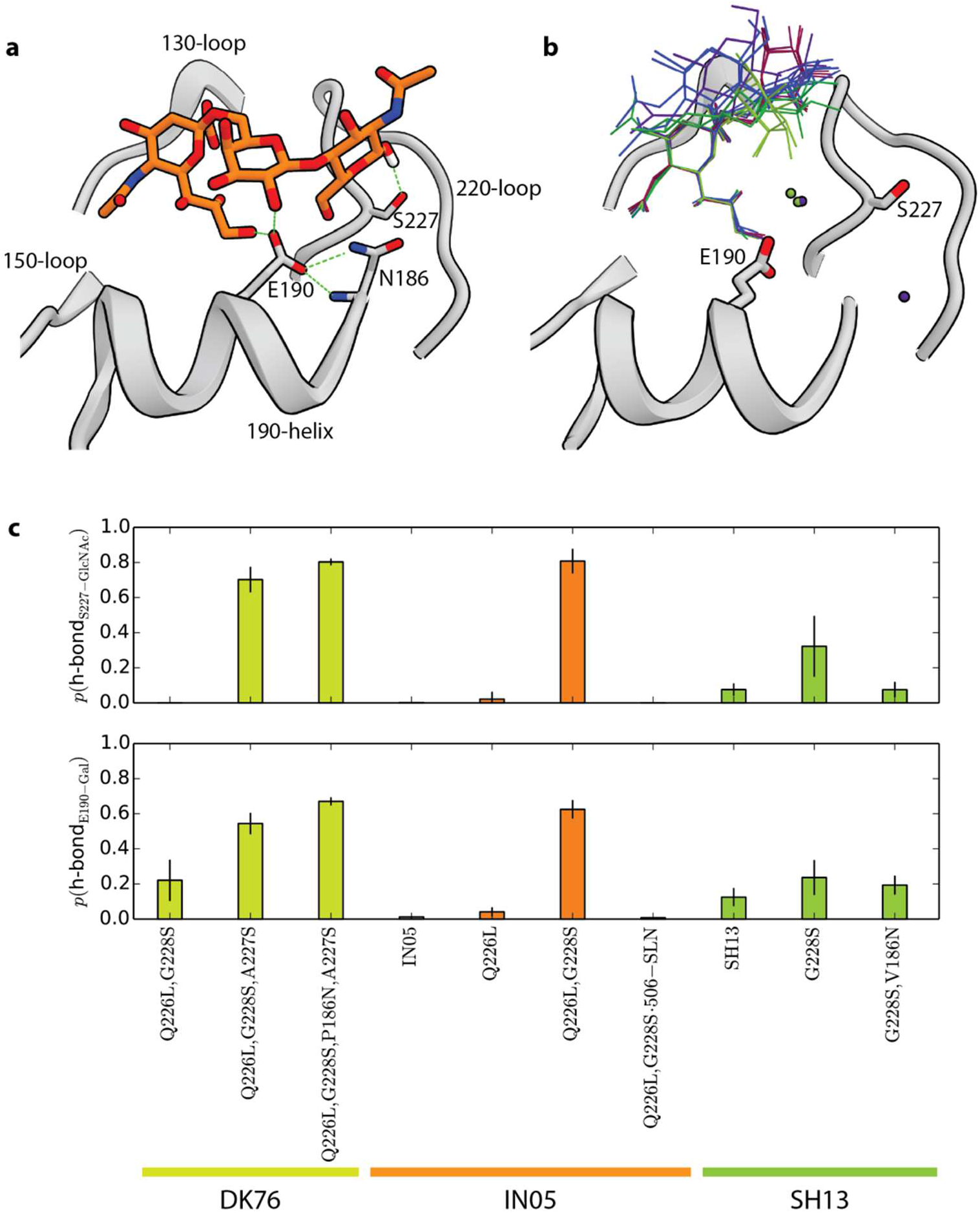
The human receptor bound to HA containing the Q226L,G228S mutations in a novel conformation not yet seen in X-ray crystallography. (**a**) A representative snapshot of the novel conformation of human receptor binding to an HA containing the Q226L,G228S mutations, taken from a simulation of 6-SLN bound to IN05^Q226L,G228S^. 6-SLN is shown in orange sticks; the characteristic hydrogen bonds in this conformation are highlighted by green dashed lines. (**b**) The structures of human receptor analogs in complex with HAs of identical or very similar sequences (PDB IDs: 4K64, 4K67, 4BGX, 4CQU, 4CQR, 4BH3, 4KDO). The spheres are water molecules present in the crystal structures. S227 does not form any direct or water-mediated hydrogen bonds with the human receptors, nor does E190 with the galactose, in any of the crystal structures: in the crystallographic structures we have examined (listed on the x-axis of Fig. S4), the minimum distance between the side chain of any residue at position 227 and the bound receptor analog is 4.2 Å, and that between the side chain of E190 and the O_2_ of galactose is 5.2 Å. (**c**) The probabilities that the hydrogen bonds between S227 and the terminal hydroxyl of GlcNac (top) and between E190 and the O_2_ hydroxyl of Gal (bottom) formed in the receptor-HA complex during our simulations.

In our MST measurements (Table 1), 6-SLN has no detectable binding to the wild-type IN05 HA, but it reproducibly bound to the Q226L,G228S mutant with *K*_*D*_ = 0.055 ± 0.004 mM, confirming that this pair of mutations increases the HA’s binding affinity for 6-SLN. When the S227A mutation is added, 6-SLN binding to HA can no longer be detected by thermophoresis, but the *K*_*D*_ value is estimated to be 1.6 ± 0.1 mM based on fluorescence quenching, a decrease in binding affinity consistent with our predictions above.

To identify new gain-of-binding mutations that exploit the novel cis/g/tg conformation, we studied DK76 HA of the H1 subtype, in which (unlike in IN05) the Q226L,G228S mutations do not improve binding to 6-SLN (Figs. 2d and 5; Table 1). In our simulations 6-SLN did, however, bind in the cis/g/tg conformation to DK76^Q226L,G228S^, though it occupied this conformation less than it did when binding to IN05^Q226L,G228S^ (Fig. 3c). We predicted that an additional pair of mutations, A227S and P186N, would establish hydrogen bonds between S227 and GlcNAc and between N186 and E190 in the cis/g/tg conformation, increasing the binding affinity of DK76^Q226L,G228S^ for 6-SLN.

Indeed, in our MD simulations, 6-SLN bound predominantly in the cis/g/tg conformation to this quadruple mutant DK76^Q226L,G228S,A227S,P186N^ (Fig. 3c), and the above two hydrogen bonds formed for the majority of the time that 6-SLN was bound (Fig. 4c). The quadruple mutant bound 6-SLN with both increased *k*_*a*_ and decreased *K*_*d*_ (Fig. 2) compared to wild-type DK76, as estimated from the simulated binding equilibrium; its estimated affinity for 6-SLN increased by a factor of 50 (Table 1). We again performed MST measurements to corroborate the results, and found that 6-SLN bound to the quadruple mutant with *K*_*D*_ = 0.06 ± 0.03 mM, compared to no detectable binding to the DK76 wild type, qualitatively consistent with the predicted affinity increase (Fig. 5; Table 1). Our simulations also predicted, and our MST experiments confirmed, that eliminating either the A227S or P186N mutation leads to a substantial reduction in affinity for 6-SLN (Table 1), suggesting that both the S227-GlcNAc and the E190-Gal hydrogen bonds contribute to the increased affinity for the human receptor.

**Figure 5.**
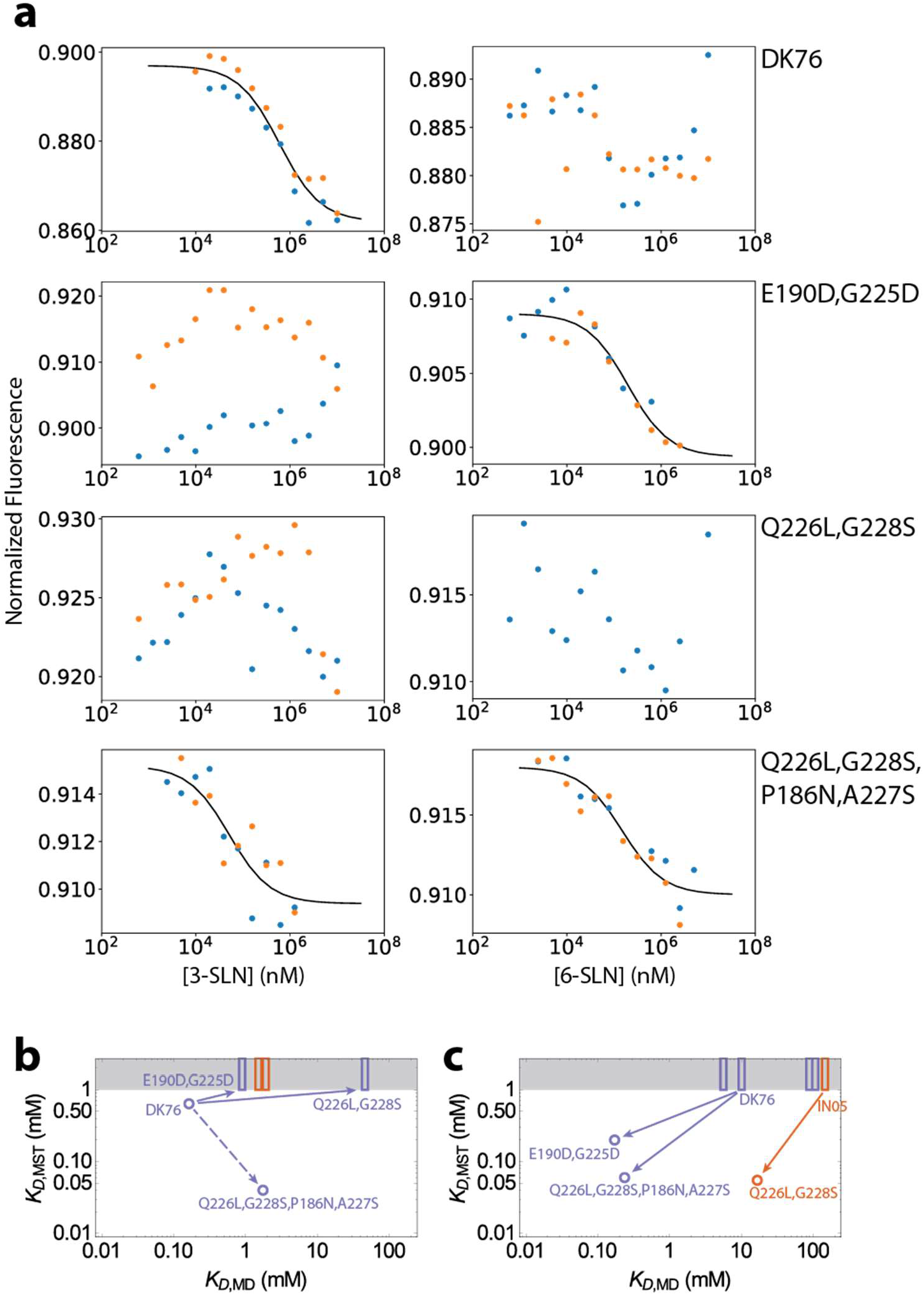
MST measurements of the binding affinities of 3-SLN and 6-SLN to selected DK76 HA variants are shown. (**a**) Titration curves for 3-SLN (left) and 6-SLN (right) binding to DK76 wild-type, DK76E190D,G225D, DK76Q226L,G228S, and DK76Q226L,G228S,P186N,A227S. In the cases of detectable binding, the normalized fluorescence decreases with the receptor analog concentration. The blue and orange points represent two replicate experiments, and the black curves are fits of the mass-action law that yield the estimated dissociation constants, *K*_*D*_, in Table 1. The experimental binding affinities are compared to the values obtained from MD simulations for 3-SLN (**b**) and 6-SLN (**c**). Binding affinities below the detection limit of MST are shown in the gray box as rectangles at their respective values estimated by MD. Binding affinities for DK76 variants are colored blue; those for IN05 variants, orange. MST and MD are in good agreement with regard to the effects of the mutations (indicated by the arrows), with the exception of the DK76 mutations Q226L,G228S,P186N,A227S on 3-SLN binding (indicated by the dashed arrow).

The different binding-conformation distributions of 6-SLN and 506-SLN (Figs. 3c and S5) suggest that HA mutations can have different effects depending on which receptor analog they bind to. 506-SLN in complex with IN05 variants visited a different set of conformations than 6-SLN did in our simulations (Fig. S5), preferred the conformation that is observed in the crystal structure of 506-SLN in complex with IN05^Q226L,G228S^,^23^ and rarely adopted the cis/g/tg conformation (Fig. 3c) or formed the hydrogen bond between E190 and the O_2_ hydroxyl of Gal (Fig. 4c). 506-SLN and complete, natural receptors, unlike the analog 6-SLN, lack the terminal hydroxyl on GlcNac and thus cannot form a hydrogen bond, as a donor, with S227. We thus predicted that the additional mutations A227S,P186N, which improved the affinity of DK76^Q226L,G228S^ for 6-SLN, would not do so in the case of 506-SLN; our MST measurements also confirmed this prediction (Table 1). Our results are consistent with the experimental observation that an HA molecule can have different avidities for different α2,6-linked glycans,^11,12,24^ and imply that different HA mutants might engage SA receptors of different glycan compositions on host cells.^6^

We note that the *K*_*D*_ values measured by our MST assays are systematically lower than previously reported values^9,25^ for HA-SA binding. There are important differences between the experiments that may account for these quantitative differences: The HA trimer concentrations, for example, are much higher in NMR experiments (~30 μM)^25^ than in MST experiments (~200 nM). At high HA and SA concentrations, nonspecific SA binding to HA may compete against the weak specific binding, thus reducing the effective SA concentration in solution and decreasing the measured binding affinity. The buffers also differ between our MST assay and the previous experiments: Most notably, Mg^2+^ is present in our MST buffer, but not in the buffers of the previous experiments, whereas phosphate is used in the buffers of the previous experiments but not in our MST buffer. Because SA contains a charged carboxylic acid in Neu5Ac and multiple polar hydroxyls, it is unsurprising that the difference in the ionic co-solvents might lead to a significant difference in the measured affinities. Furthermore, our MST measurements used the label-free method, which avoids the non-specific labeling of HA that can potentially interfere with SA binding and result in a lower measured affinity. Despite the quantitative differences in the absolute *K*_*D*_ values for previously characterized mutations,^9^ the results agree in the directional changes in the binding affinities, which are more relevant than absolute *K*_*D*_ values for identifying affinity-switching mutations. We thus believe that our predictions of the affinity-switching mutations will hold true under different experimental conditions.

Also worth noting are the quantitative differences between the MD-estimated and the MST-measured *K*_*D*_ values. Although our MD simulations were long enough for some of the HA-SA pairs to yield relatively precise estimates of *K*_*D*_ values, they were still too short for pairs with slow binding or unbinding kinetics to have numbers of binding and unbinding events sufficient to produce such estimates. The 95% confidence intervals for the estimated *K*_*D*_ values of 3-SLN binding to DK76 (0.01‒0.7 mM) and to DK76^Q226L,G228S,P186N,A227S^ (0.3‒14.2 mM), for example, overlap, which can confound the prediction of the direction of affinity change. Furthermore, our MD simulations modeled the binding of SA to an HA RBD monomer, whereas MST measured the binding of SA to the HA trimer, implying that MD would systematically underestimate *K*_*D*_ by a factor of 3 compared to MST. In addition, the co-solvents in the MST buffer are not included in the simulations (a common limitation for MD simulations, as many of the co-solvents and bivalent ions do not have well-parameterized force field models), which, as discussed above, could contribute a systematic difference in calculated *K*_*D*_ values. Insufficient accuracy of the force field may introduce additional deviations. Nonetheless, the affinity-switching mutations identified by the MD-predicted *K*_*D*_ changes are largely in agreement with the MST measurements. This finding suggests that long timescale MD simulations can be used not only to generate SA-binding conformations for prediction of potential human-adaptive HA mutations, but also to provide an initial assessment of the effect of such mutations on the HA-SA binding affinities.

A notable feature of the simulations we report is their timescale, which is orders-of-magnitude longer than previous simulations of HA-SA complexes.^27,37^ Due to their length, our simulations are able to generate SA binding conformations not previously reported in MD simulations or in crystallography, and to give statistically converged estimates of the relative populations of SA binding conformations to a wide range of HA variants, including H1, H5, and H7 subtypes. This represent new structural and thermodynamic information that can be used in structure-based prediction of potential affinity-switching mutations, as illustrated in this work. In addition, our long-timescale MD simulations enabled the unbiased simulations of SA binding to and unbinding from HA, yielding estimates of the corresponding kinetic rate constants, which are difficult to measure experimentally for these weak complexes.

Taken together, our simulations suggest that a human receptor binds to a given avian HA in a diverse ensemble of conformations, and that specificity-switching mutations induce new favorable interactions in one of these conformations, stabilizing that conformation and promoting affinity for the human receptor. The conformational diversity of the bound receptors increases the number of viable mutations, potentially facilitating the human adaptation of the influenza virus. The conformations generated by our MD simulations may potentially serve as starting points for further analyses,^27,37^ such as free energy calculations^29^ and for rational, structure-based protein design,^38^ to help identify influenza strains on the verge of human transmission.

## Methods Summary

### Molecular dynamics simulations

The starting structures of the receptor-binding domains (RBDs) of the HA molecules were taken from the corresponding crystal structures (DK76 from PDB ID: 2WRH; IN05 from PDB ID: 4K64; and SH13 from PDB ID: 4LKG); the mutations corresponding to different variants were introduced and modeled using the Maestro software.^39^ The RBD of DK76 in our simulation and in the crystal structure 2WRH differ from the RBD of A/duck/Alberta/35/76 (GenBank accession number: AF091309) by a deletion of threonine at position 132, outside the binding pocket. Comparison with a closely related sequence with the additional threonine at position 132, in the PDB structure 3HTQ, suggests that the additional threonine creates a bulge outside the binding pocket and does not impact the pocket’s structure. Structural models of the receptor analogs (3-SLN, 6-SLN, and 506-SLN) were built using the Glycam web server.^40^ The Amber99SB-ILDN force field^41,42^ was used for the proteins, the Glycam06^43^ force field was used for the polysaccharides (3-SLN, 6-SLN, and 506-SLN), and the TIP3P water model was used. Isobaric-isothermal simulations, as described previously,^44^ were run at a temperature of 310 K and a pressure of 1 atm, using the Anton specialized hardware.^31^

### Expression and purification of HA proteins and MST

Trimers of HA variants were produced, and binding to 3-SLN, 6-SLN, and 506-SLN was measured, using MST by Crelux (now part of Wuxi Apptec) through contracted research. (The experimental details are given in SI.) To eliminate bias in data interpretation, Crelux was not informed of the predictions of the MD simulations at any point in the course of the research.

## Acknowledgments

We thank Qi Wang, David Borhani, Robert McGibbon, and Michael Eastwood for helpful discussions and critical readings of the manuscript, Caleb Jordan and Kevin Yuh for preparing the molecular graphics, Ansgar Philippsen for preparing the supplementary movie, and Rebecca Bish-Cornelissen and Berkman Frank for editorial assistance.

## References

1. Kuiken T, Holmes EC, McCauley J, Rimmelzwaan GF, Williams CS, Grenfell BT. Host species barriers to influenza virus infections. Science 312(5772), 394–397 (2006).

2. Morens DM, Subbarao K, Taubenberger JK. Engineering H5N1 avian influenza viruses to study human adaptation. Nature 486(7403), 335–340 (2012).

3. Baigent SJ, McCauley JW. Influenza type A in humans, mammals and birds: determinants of virus virulence, host-range and interspecies transmission. Bioessays 25(7), 657–671 (2003).

4. Skehel JJ, Wiley DC. Receptor binding and membrane fusion in virus entry: the influenza hemagglutinin. Annu. Rev. Biochem. 69, 531–569 (2000).

5. Steinhauer DA. Influenza: Pathways to human adaptation. Nature 499(7459), 412–413 (2013).

6. Xu H, Shaw DE. A Simple Model of Multivalent Adhesion and Its Application to Influenza Infection. Biophys. J. 110(1), 218–233 (2016).

7. Couceiro JN, Paulson JC, Baum LG. Influenza virus strains selectively recognize sialyloligosaccharides on human respiratory epithelium; the role of the host cell in selection of hemagglutinin receptor specificity. Virus Res. 29(2), 155–165 (1993).

8. Stevens J, Blixt O, Tumpey TM, Taubenberger JK, Paulson JC, Wilson IA. Structure and receptor specificity of the hemagglutinin from an H5N1 influenza virus. Science 312(5772), 404–410 (2006).

9. Xiong X, Coombs PJ, Martin SR, Liu J, Xiao H, McCauley JW, Locher K, Walker PA, Collins PJ, Kawaoka Y, Skehel JJ, Gamblin SJ. Receptor binding by a ferret-transmissible H5 avian influenza virus. Nature 497(7449), 392–396 (2013).

10. Tharakaraman K, Jayaraman A, Raman R, Viswanathan K, Stebbins NW, Johnson D, Shriver Z, Sasisekharan V, Sasisekharan R. Glycan receptor binding of the influenza A virus H7N9 hemagglutinin. Cell 153(7), 1486–1493 (2013).

11. Watanabe T, Kiso M, Fukuyama S, Nakajima N, Imai M, Yamada S, Murakami S, Yamayoshi S, Iwatsuki-Horimoto K, Sakoda Y, Takashita E, McBride R, Noda T, Hatta M, Imai H, Zhao D, Kishida N, Shirakura M, de Vries RP, Shichinohe S, Okamatsu M, Tamura T, Tomita Y, Fujimoto N, Goto K, Katsura H, Kawakami E, Ishikawa I, Watanabe S, Ito M, Sakai-Tagawa Y, Sugita Y, Uraki R, Yamaji R, Eisfeld AJ, Zhong G, Fan S, Ping J, Maher EA, Hanson A, Uchida Y, Saito T, Ozawa M, Neumann G, Kida H, Odagiri T, Paulson JC, Hasegawa H, Tashiro M, Kawaoka Y. Characterization of H7N9 influenza A viruses isolated from humans. Nature 501(7468), 551–555 (2013).

12. Chandrasekaran A, Srinivasan A, Raman R, Viswanathan K, Raguram S, Tumpey TM, Sasisekharan V, Sasisekharan R. Glycan topology determines human adaptation of avian H5N1 virus hemagglutinin. Nat. Biotechnol. 26(1), 107–113 (2008).

13. Rogers GN, Daniels RS, Skehel JJ, Wiley DC, Wang XF, Higa HH, Paulson JC. Host-mediated selection of influenza virus receptor variants. Sialic acid-alpha 2,6Gal-specific clones of A/duck/Ukraine/1/63 revert to sialic acid-alpha 2,3Gal-specific wild type in ovo. J. Biol. Chem. 260(12), 7362–7367 (1985).

14. Imai M, Kawaoka Y. The role of receptor binding specificity in interspecies transmission of influenza viruses. Curr. Opin. Virol. 2(2), (2012).

15. Matrosovich MN, Gambaryan AS, Teneberg S, Piskarev VE, Yamnikova SS, Lvov DK, Robertson JS, Karlsson KA. Avian influenza A viruses differ from human viruses by recognition of sialyloligosaccharides and gangliosides and by a higher conservation of the HA receptor-binding site. Virology 233(1), 224–234 (1997).

16. Air, GM. Sequence relationships among the hemagglutinin genes of 12 subtypes of influenza A virus. Proc. Natl. Acad. Sci. U.S.A. 78(12), 7639–7643 (1981).

17. Matrosovich M, Tuzikov A, Bovin N, Gambaryan A, Klimov A, Castrucci MR, Donatelli I, Kawaoka Y. Early alterations of the receptor-binding properties of H1, H2, and H3 avian influenza virus hemagglutinins after their introduction into mammals. J. Virol. 74(18), 8502–8512 (2000).

18. Kasson PM, Ensign DL, Pande VS. Combining molecular dynamics with Bayesian analysis to predict and evaluate ligand-binding mutations in influenza hemagglutinin. J. Am. Chem. Soc. 101(32), 11338–11340 (2009).

19. Maurer-Stroh S, Li Y, Bastien N, Gunalan V, Lee RT, Eisenhaber F, Booth TF. Potential human adaptation mutation of influenza A(H5N1) virus, Canada. Emerg. Infect. Dis. 20(9), 1580–1582 (2014).

20. Jongkon N, Mokmak W, Chuakheaw D, Shaw PJ, Tongsima S, Sangma C. Prediction of avian influenza A binding preference to human receptor using conformational analysis of receptor bound to hemagglutinin. BMC Genomics 10 (Suppl. 3), S24 (2009).

21. Liu J, Stevens DJ, Haire LF, Walker PA, Coombs PJ, Russell RJ, Gamblin SJ, Skehel JJ. Structures of receptor complexes formed by hemagglutinins from the Asian Influenza pandemic of 1957. Proc. Natl. Acad. Sci. U.S.A. 106(40), 17175–17180 (2009).

22. Shi Y, Zhang W, Wang F, Qi J, Wu Y, Song H, Gao F, Bi Y, Zhang Y, Fan Z, Qin C, Sun H, Liu J, Haywood J, Liu W, Gong W, Wang D, Shu Y, Wang Y, Yan J, Gao GF. Structures and receptor binding of hemagglutinins from human-infecting H7N9 influenza viruses. Science 342(6155), 243–247 (2013).

23. Zhang W, Shi Y, Lu X, Shu Y, Qi J, Gao GF. An airborne transmissible avian influenza H5 hemagglutinin seen at the atomic level. Science 340(6139), 1463–1467 (2013).

24. de Vries RP, Zhu X, McBride R, Rigter A, Hanson A, Zhong G, Hatta M, Xu R, Yu W, Kawaoka Y, de Haan CA, Wilson IA, Paulson JC. Hemagglutinin receptor specificity and structural analyses of respiratory droplet-transmissible H5N1 viruses. J. Virol. 88(1), 768–773 (2014).

25. Sauter NK, Bednarski MD, Wurzburg BA, Hanson JE, Whitesides GM, Skehel JJ, Wiley DC. Hemagglutinins from two influenza virus variants bind to sialic acid derivatives with millimolar dissociation constants: a 500-MHz proton nuclear magnetic resonance study. Biochemistry 28(21), 8388–8396 (1989).

26. Vachieri SG, Xiong X, Collins PJ, Walker PA, Martin SR, Haire LF, Zhang Y, McCauley JW, Gamblin SJ, Skehel JJ. Nature 511(7510), 475–477 (2014).

27. Elli S, Macchi E, Rudd TR, Raman R, Sassaki G, Viswanathan K, Yates EA, Shriver Z, Naggi A, Torri G, Sasisekharan R, Guerrini M. Insights into the human glycan receptor conformation of 1918 pandemic hemagglutinin-glycan complexes derived from nuclear magnetic resonance and molecular dynamics studies. Biochemistry 53(25), 4122–4135 (2014).

28. Gamblin SJ, Haire LF, Russell RJ, Stevens DJ, Xiao B, Ha Y, Vasisht N, Steinhauer DA, Daniels RS, Elliot A, Wiley DC, Skehel JJ. The structure and receptor binding properties of the 1918 influenza hemagglutinin. Science 303(5665), 1838–1842 (2004).

29. Das P, Li J, Royyuru AK, Zhou R. Free energy simulations reveal a double mutant avian H5N1 virus hemagglutinin with altered receptor binding specificity. J. Comput. Chem. 30(11), 1654–1663 (2009).

30. Dror RO, Dirks RM, Grossman JP, Xu H, Shaw DE. Biomolecular simulation: a computational microscope for molecular biology. Annu. Rev. Biophys. 41, 429–452 (2012).

31. Shaw DE, Grossman JP, Bank JA, Batson B, Butts JA, Chao JC, Deneroff MM, Dror RO, Even A, Fenton CH, Forte A, Gagliardo J, Gill G, Greskamp B, Ho CR, Ierardi DJ, Iserovich L, Kuskin JS, Larson RH, Layman T, Lee L-S, Lerer AK, Li C, Killebrew D, Mackenzie KM, Mok SY-H, Moraes MA, Mueller R, Nociolo LJ, Peticolas JL, Quan T, Ramot D, Salmon JK, Scarpazza DP, Schafer UB, Siddique N, Snyder CW, Spengler J, Tang PTP, Theobald M, Toma H, Towles B, Vitale B, Wang SC, Young C. Anton 2: Raising the bar for performance and programmability in a special-purpose molecular dynamics supercomputer. Proceedings of the International Conference for High Performance Computing, Networking, Storage and Analysis (SC14) New York, NY: IEEE (2014).

32. Imai M, Watanabe T, Hatta M, Das SC, Ozawa M, Shinya K, Zhong G, Hanson A, Katsura H, Watanabe S, Li C, Kawakami E, Yamada S, Kiso M, Suzuki Y, Maher EA, Neumann G, Kawaoka Y. Experimental adaptation of an influenza H5 HA confers respiratory droplet transmission to a reassortant H5 HA/H1N1 virus in ferrets. Nature 486(7403), 420–428 (2012).

33. Herfst S, Schrauwen EJ, Linster M, Chutinimitkul S, de Wit E, Munster VJ, Sorrell EM, Bestebroer TM, Burke DF, Smith DJ, Rimmelzwaan GF, Osterhaus AD, Fouchier RA. Airborne transmission of influenza A/H5N1 virus between ferrets. Science 336(6088), 1534–1541 (2012).

34. Tumpey TM, Maines TR, Van Hoeven N, Glaser L, Solórzano A, Pappas C, Cox NJ, Swayne DE, Palese P, Katz JM, García-Sastre A. A two-amino acid change in the hemagglutinin of the 1918 influenza virus abolishes transmission. Science 315(5812), 655–659 (2007).

35. Mobley DL, Dill KA. Binding of small-molecule ligands to proteins: “what you see” is not always “what you get.” Structure 17(4), 489–498 (2009).

36. Wienken CJ, Baaske P, Rothbauer U, Braun D, Duhr S. Protein-binding assays in biological liquids using microscale thermophoresis. Nat. Commun. 1, 100 (2010).

37. Xu D, Newhouse EI, Amaro RE, Pao HC, Cheng LS, Markwick PR, McCammon JA, Li WW, Arzberger PW. Distinct glycan topology for avian and human sialopentasaccharide receptor analogues upon binding different hemagglutinins: a molecular dynamics perspective. J. Mol. Biol. 387(2), 465–491 (2009).

38. Tinberg CE, Khare SD, Dou J, Doyle L, Nelson JW, Schena A, Jankowski W, Kalodimos CG, Johnsson K, Stoddard BL, Baker D. Computational design of ligand-binding proteins with high affinity and selectivity. Nature 501(7466), 212–216 (2013).

39. Schrödinger Release 2017-3: Maestro, Schrödinger, LLC, New York, NY, 2017. (http://www.schrodinger.com/Maestro)

40. Woods Group. (2005–2017) GLYCAM Web. Complex Carbohydrate Research Center, University of Georgia, Athens, GA. (http://glycam.org)

41. Hornak V, Abel R, Okur A, Strockbine B, Roitberg A, Simmerling C. Comparison of multiple Amber force fields and development of improved protein backbone parameters. Proteins 65(3), 712–725 (2006).

42. Lindorff-Larsen K, Piana S, Palmo K, Maragakis P, Klepeis JL, Dror RO, Shaw DE. Improved side-chain torsion potentials for the Amber ff99SB protein force field. Proteins 78(8), 1950–1958 (2010).

43. Kirschner KN, Yongye AB, Tschampel SM, González-Outeiriño J, Daniels CR, Foley BL, Woods RJ. GLYCAM06: a generalizable biomolecular force field. Carbohydrates. J. Comput. Chem. 29(4), 622–655 (2008).

44. Schmidt AG, Xu H, Khan AR, O'Donnell T, Khurana S, King LR, Manischewitz J, Golding H, Suphaphiphat P, Carfi A, Settembre EC, Dormitzer PR, Kepler TB, Zhang R, Moody MA, Haynes BF, Liao HX, Shaw DE, Harrison SC. Preconfiguration of the antigen-binding site during affinity maturation of a broadly neutralizing influenza virus antibody. Proc. Natl. Acad. Sci. U.S.A. 110(1), 264–269 (2013).

45. Poppe L, Stuike-Prill R, Meyer B, van Halbeek H. The solution conformation of sialyl-alpha (2→6)-lactose studied by modern NMR techniques and Monte Carlo simulations. J. Biomol. NMR 2(2), 109–136 (1992).

46. Sassaki GL, Elli S, Rudd TR, Macchi E, Yates EA, Naggi A, Shriver Z, Raman R, Sasisekharan R, Torri G, Guerrini M. Human (α2→6) and avian (α2→3) sialylated receptors of influenza A virus show distinct conformations and dynamics in solution. Biochemistry 52(41), 7217–7230 (2013).

