## Supplementary Information for "Characterizing receptor flexibility to predict mutations that lead to human adaptation of influenza hemagglutinin"

### Supplementary Methods

#### *Estimation of the kinetic rates of binding and unbinding from the simulations*

We treat binding and unbinding as Poisson processes. For the binding process, the rate constant of the Poisson process is $k = k_a[L]$, where $k_a$ is the association rate constant, and $[L]$ is the receptor analog concentration in the corresponding simulations; for the unbinding process, the rate constant of the Poisson process is the dissociation rate constant $k_d$.

Given $N$ independent instances of the same Poisson process where a random event occurs at the constant rate $k$, the probability that $n$ events have occurred at times $\{t_{i=1,2,\ldots,n}\}$, and that no event has occurred in the other $m = N - n$ attempts that terminated after times $\{T_{j=1,2,\ldots,m}\}$, is

$$p(\{t_i\},\{T_j\}|k) = \prod_{i=1}^{n} k \exp(-kt_i) \prod_{j=1}^{m} \exp(-kT_j) = k^n \exp\left(-k\left(\sum_{i=1}^{n} t_i + \sum_{j=1}^{m} T_j\right)\right)$$

$$\equiv k^n \exp(-kt),$$

where we have introduced the characteristic time $t \equiv \sum_{i=1}^{n} t_i + \sum_{j=1}^{m} T_j$. This probability thus only depends on $n$ and $t$: $p(\{t_i\},\{T_j\}|k) = p(n,t|k)$. The maximum likelihood estimator (MLE) of $k$ is:

$$k = n / t.$$

The variance on the estimate is given by:

$$\sigma_k^2 = -\left(\frac{\partial^2 \ln p}{\partial k^2}\right)^{-1} = \frac{k^2}{n}$$

or

$$\sigma_{\ln k}^2 = n^{-1}$$

We report the above MLE of $k$ and its associated variance when $n > 0$, (i.e., we report the MLE of $k_a$ when there are binding events in the simulations, and the MLE of $k_d$ when there are unbinding events in the simulations).

In the case that no event occurs (i.e., $n = 0$), we have $\sigma_k = \infty$, and we cannot have a meaningful MLE for the value of $k$. Nevertheless, we can estimate an upper bound of $k$ given the terminal times $\{T_{j=1,2,\ldots,m}\}$, as described below. The posterior probability of $k$, given the number of events $n$ and the characteristic time $t$, is

$$p(k|n,t) = \frac{p(n,t|k)p(k)}{\int dk' p(n,t|k')p(k')}.$$

Choosing the improper uniform prior $p(k) = 1 / K$, for $0 < k \leq K$, we have

$$p(k|n,t) = \lim_{K\to\infty} \frac{\frac{p(n,t|k)}{K}}{\int_0^K \frac{dk' p(n,t|k')}{K}} = \frac{k^n \exp(-kt)}{n! t^{-1-n}}.$$

If $n = 0$, the probability that $k > k_u$ is

$$p(k > k_u|n,t) = \int_{k_u}^{\infty} dk' p(k'|n,t) = \exp(-k_u t).$$

Given a value $P \in (0,1)$, we can derive an upper bound $k_u$, such that the probability that the true rate constant is greater than the upper bound is $p(k > k_u|n,t) = P$, to be

$$k_u = -\ln P / t.$$

***Estimation of the equilibrium dissociation constant from simulated binding equilibrium***

If we have both binding and unbinding events, we can estimate both values of $k_a$ and $k_d$, and thus the value of $K_D = k_d / k_a$.

If we have only binding events but no unbinding events, we can estimate the value of $k_a$ and establish an upper bound of $k_d$ (see above), and thus an upper bound of $K_D$. Suppose that in simulating binding, we have $n_a$ successful binding events at times $\{t_{a,i=1,\ldots,n_a}\}$ and $m_a$ unsuccessful attempts terminating after times $\{T_{a,j=1,\ldots,m_a}\}$, and in simulating unbinding, we have $n_d$ successful unbinding events at times $\{t_{d,i=1,\ldots,n_d}\}$ and $m_d$ unsuccessful attempts terminating after times $\{T_{d,j=1,\ldots,m_d}\}$. Let $t_s = \sum_{i=1}^{n_s} t_{s,i} + \sum_{j=1}^{m_s} T_{s,j}$ to be the characteristic times for the binding ($s = a$) and the unbinding ($s = d$) processes. The posterior probability that the association rate has the value $k_a$ and the dissociation rate has the value $k_d$ is

$$p(k_a, k_d|n_a, t_a, n_d, t_d) = p(k_a|n_a, t_a)p(k_d|n_d, t_d)$$

Given a value $K_D$, the probability that $k_d / k_a \geq K_D$ is

$$p\left(\frac{k_d}{k_a} \geq K_D|n_a, t_a, n_d = 0, t_d\right)$$

$$= \iint_{\frac{k_d}{k_a} \geq K_D} dk_a dk_d p(k_a, k_d|n_a, t_a, n_d = 0, t_d)$$

$$= \int_0^\infty dk_a \frac{k_a^{n_a} \exp(-k_a t_a)}{n_a! t_a^{-1-n_a}} \int_{k_a K_D}^\infty dk_d t_d \exp(-k_d t_d) = \left(\frac{t_a}{t_a + K_D t_d}\right)^{n_a+1}.$$

Given a value $P \in (0,1)$, we can establish an upper bound, $K_D$, of the equilibrium constant, such that the probability that the true equilibrium constant is greater than the upper bound is $p(k_d / k_a \geq K_D) = P$, to be

$$K_D = \frac{t_a}{t_d}\left(P^{-1/(n_a+1)} - 1\right)$$

If there are a large number of binding events (i.e., $n_a >> 1$), the estimate of $k_a$ becomes very precise, and the upper bound of $K_D$ approaches the ratio between $k_a$ and the upper bound of $k_d$, since

$$\lim_{n_a \to \infty} \frac{t_a}{t_d}\left(P^{-1/(n_a+1)} - 1\right) = \lim_{n_a \to \infty} \frac{t_a/n_a}{t_d}\left(n_a\left(e^{-\frac{1}{n_a+1}\ln P} - 1\right)\right) = \frac{-\ln P / t_d}{k_a}$$

Conversely, if we have unbinding events but no binding events, we can establish a lower bound, $K_D$, of the equilibrium constant for a given value $P \in (0,1)$, such that

$$P = p\left(\frac{k_d}{k_a} \leq K_D | n_a = 0, t_a, n_d, t_d\right) = \left(\frac{t_d}{t_d + K_D^{-1} t_a}\right)^{n_d+1}$$

to be

$$K_D = \frac{t_a}{t_d}\left(P^{-1/(n_d+1)} - 1\right)^{-1}$$

#### *Expression of hemagglutinin variants*

SF21 cell stocks were maintained below $8 \times 10^{16}$ cells / mL and monitored for cell diameters to be around 12 µm and culture vitalities to be >95%. The production stocks were exchanged every month. Hemagglutinin (HA) was overproduced in 3 L Erlenmeyer cell culture flasks (Corning).

HA-expressing recombinant baculoviruses were generated using the flashBAC ULTRA™ system (Oxford Expression Technologies) according to the manufacturer's protocol. Multiples of 0.75 L suspension cultures with $2 \times 10^6$ SF21 cells / mL in Insect XPRESS™ (Lonza) were synchronized for 5 hours and afterwards transfected with 2.7 mL recombinant virus stock. The expressed proteins carried a C-terminal extension: SGR*ENLYFQG*G<u>GSGYIPEAPRDGQAYVRKDGEWVLLSTFL</u>GHHHHHH, where the italicized sequence is the recognition site of the TEV protease cleavage, and the underlined sequence is a trimerization foldon.[1] Infected cultures were incubated at 27 °C and 120 rpm. After 66 hours the cultures were monitored for vitality, cell diameter, and cell number. Cultures were harvested by centrifugation at 500 g for 40 minutes at 4 °C. Resulting supernatants were immediately used for purification.

***Purification of HA variants***

400 mL of the clarified supernatant was supplemented with protease inhibitors, titrated to pH 7.0 using Bis-Tris-Propane, and incubated with 5 mL Ni-NTA (HisPur, Thermo Fisher Scientific) overnight on a rotating wheel. Following incubation, the Ni-NTA column was washed with 5 times the column volume of 50 mM $NaH_2PO_4$, 300 mM NaCl, 20 mM Imidazole, 0.1% Tween 20, at pH 8.0, and eluted with 2 times the column volume of 50 mM $NaH_2PO_4$, 300 mM NaCl, 500 mM Imidazole, 0.1% Tween 20, at pH 8.0. The HA-containing fractions were determined by SDS-PAGE, pooled, and further purified to homogeneity by size-exclusion chromatography using an S200 column equilibrated with 20 mM Tris-HCl, 150 mM NaCl, at pH 7.5. ~6 mg of HA could be purified from 1 L of supernatant.

***Microscale thermophoresis***

Microscale thermophoresis (MST) measurements were performed on a NanoTemper Monolith NT.Labelfree instrument (NanoTemper Technologies GmbH), using the label-free method (detecting the fluorescence of the tryptophans in HA). Sodium salts of 3-SLN, 6-SLN, or 506-SLN (Dextra) were titrated in a 1:1 serial dilution series from a maximum concentration of 5–10 mM to produce a 12–16 point measurement in either 50 mM HEPES, 100 mM NaCl, pH 7.5, or 20 mM Tris, 170 mM NaCl, 10 mM $MgCl_2$, 0.02% Tween, pH 8.0 buffer solutions, containing various HA constructs at the trimer concentration of 123, 133, or 217 nM. These titration series were then loaded into NT.LabelFree zero background standard glass capillaries, and MST measurements were made at 23 °C using 20% light-emitting-diode power and varying infrared-laser power (20–80%), depending on the construct characteristics. The laser-on time was 25 seconds, and the laser-off time was 5 seconds.

### Supplementary References


1. Stevens J, Corper AL, Basler CF, Taubenberger JK, Palese P, Wilson IA. Structure of the uncleaved human H1 hemagglutinin from the extinct 1918 influenza virus. *Science* **303(5665)**, 1866–1870 (2004).

2. Zhang W, Shi Y, Lu X, Shu Y, Qi J, Gao GF. An airborne transmissible avian influenza H5 hemagglutinin seen at the atomic level. *Science* **340(6139)**, 1463–1467 (2013).

3. Xiong X, Coombs PJ, Martin SR, Liu J, Xiao H, McCauley JW, Locher K, Walker PA, Collins PJ, Kawaoka Y, Skehel JJ, Gamblin SJ. Receptor binding by a ferret-transmissible H5 avian influenza virus. Nature 497(7449), 392–396 (2013).

4. These may also be attributable to the plasticity in bound conformations rather than difference between 6-SLN and 506-SLN.

5. Xu R, de Vries RP, Zhu X, Nycholat CM, McBride R, Yu W, Paulson JC, Wilson IA. Preferential recognition of avian-like receptors in human influenza A H7N9 viruses. *Science* **342(6163)**, 1230–1235 (2013).

6. Poppe L, Stuike-Prill R, Meyer B, van Halbeek H. The solution conformation of sialyl-alpha (2→6)-lactose studied by modern NMR techniques and Monte Carlo simulations. *J. Biomol. NMR* **2(2)**, 109–136 (1992).

7. Haasnoot CAG, de Leeuw FAAM, Altona C. The relationship between proton-proton NMR coupling constants and substituent electronegativities—I: An empirical generalization of the karplus equation. *Tetrahedron* **36(19)**, 2783–2792 (1980).

8. Spoormaker T, de Bie MJA. Empirical determination of the torsion angle dependence of the vicinal $^{13}$C-H coupling constant in a $^{13}CH_3$-C($R^1$)OH-C($R^2$)H-*H fragment of model compounds. *Recueil des Travaux Chimiques des Pays-Bas* **97(3)**, 85–87 (1978).

9. Tvaroska I, Hricovíni M, Petráková E. An attempt to derive a new Karplus-type equation of vicinal proton-carbon coupling constants for COCH segments of bonded atoms. *Carbohydrate Res.* **189**, 359–362 (1989).

10. Sassaki GL, Elli S, Rudd TR, Macchi E, Yates EA, Naggi A, Shriver Z, Raman R, Sasisekharan R, Torri G, Guerrini M. Human (α2→6) and avian (α2→3) sialylated receptors of influenza A virus show distinct conformations and dynamics in solution. *Biochemistry* **52(41)**, 7217–7230 (2013).

### Supplementary Figures and Tables

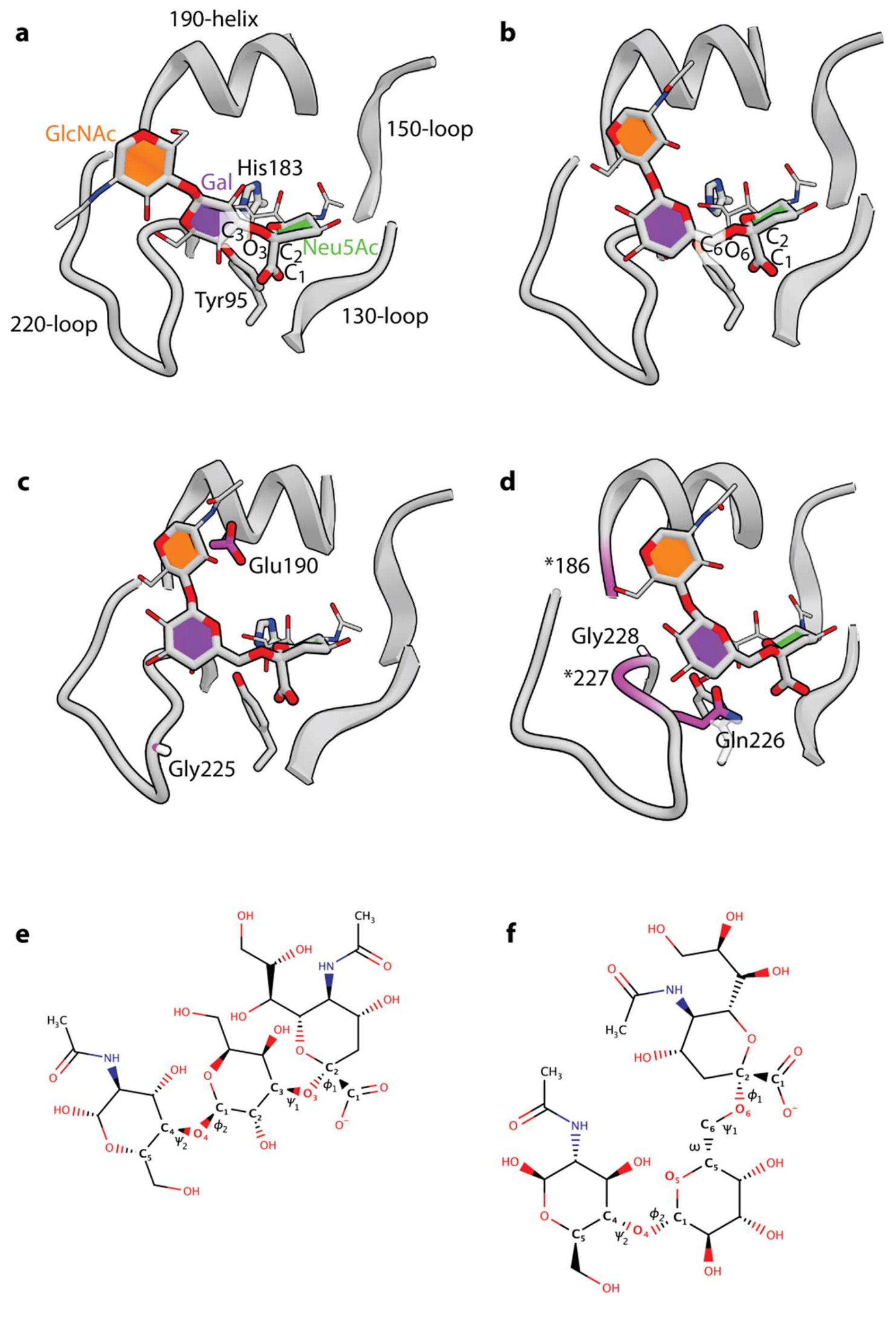


**Figure S1**. Schematics of the structure of the HA binding pocket in complex with the receptor analogs. (**a**) α2,3-linked and (**b**) α2,6-linked sialic acid (SA) receptors bound to HA in the binding pocket formed by the 130-loop, the 150-loop, the 190-helix, and the 220-loop. The binding conformations are single snapshots taken from MD simulations of 3-SLN and 6-SLN in complex with DK76. Only the linked sugar rings and the carboxylate are shown for the receptors. The two conserved residues Tyr95 and His183 are part of the binding pocket. (**c**) The residues Glu190 and Gly225 are mutated in the pair of switching mutations E190D,G225D. (**d**) The residues Gln226 and Gly228 are mutated in the pair of switching mutations Q226L,G228S. Both pairs of mutations primarily alter HA's interactions with Gal and GlyNac moieties. In (d), the purple color in the chain highlights residue positions 227 and 186, which are mutated in the quadruple mutant DK76$^{Q226L,G228S,P186N,G227S}$, enabling the H1 HA DK76 to favor binding to the human receptor analog 6-SLN over the avian receptor analog 3-SLN. The receptor analogs 3-SLN (**e**) and 6-SLN (**f**) are used in this work. The torsions used to characterize the receptor binding conformations, $(\phi_1, \psi_1, \varpi, \phi_2, \psi_2)$, are labeled.

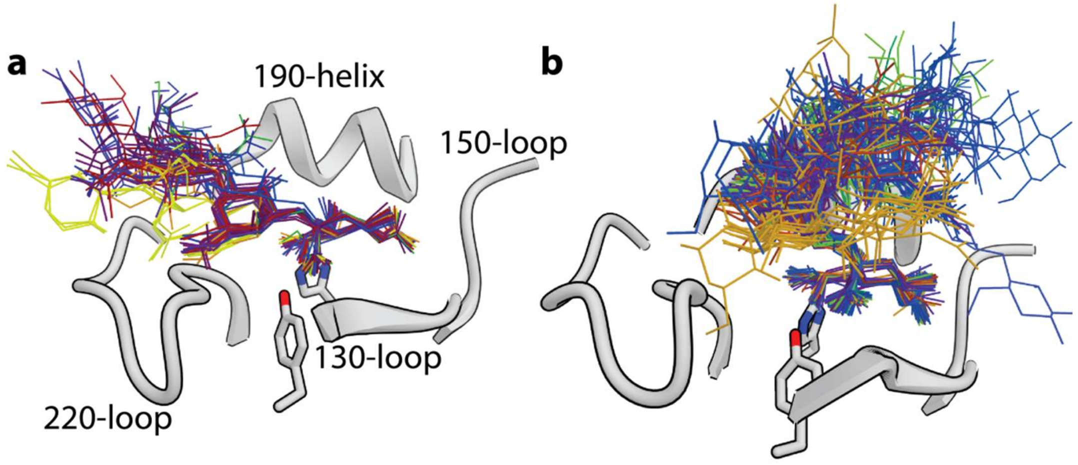


**Figure S2.** Receptor analogs in complex with HA, adopting a variety of binding conformations. In complex structures determined by X-ray crystallography for different HA variants, (**a**) α2,3-linked SAs by and large exit the binding pocket over the 220-loop, whereas (**b**) α2,6-linked SAs adopt more diverse binding conformations, exiting the binding pocket by way of a wide range of angles. The receptor analogs are shown as thin sticks, with similar structures having similar colors.

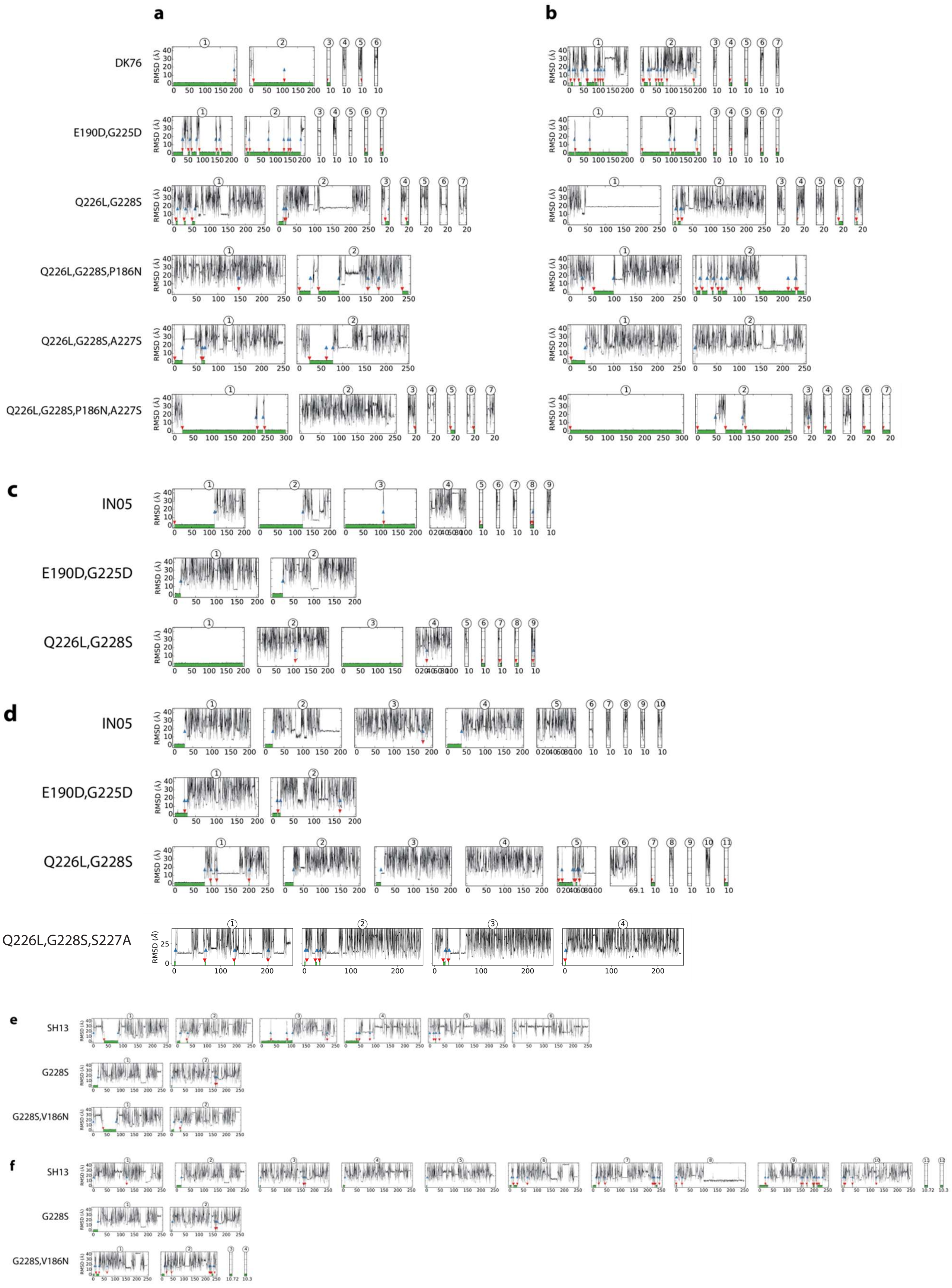


**Figure S3.** Time traces of the root-mean-square deviation (RMSD) of Neu5Ac (relative to its bound pose in crystal structures) in the simulations of 3-SLN and 6-SLN binding to the receptor-binding domain of various HAs. (**a**) Simulations of 3-SLN binding to DK76 HA variants. (**b**) Simulations of 6-SLN binding to DK76 HA variants. (**c**) Simulations of 3-SLN binding to IN05 HA variants. (**d**) Simulations of 6-SLN binding to IN05 HA variants. (**e**) Simulations of 3-SLN binding to SH13 HA variants. (**f**) Simulations of 6-SLN binding to SH13 HA variants. Individual simulations are indicated by circled numbers on top of each plot. The times of binding (when RMSD fell below 1.5 Å) and unbinding (when RMSD rose above 8 Å) are indicated by red and blue arrows, respectively, and the periods when the receptor analogs were bound are indicated by green shading under the curves.

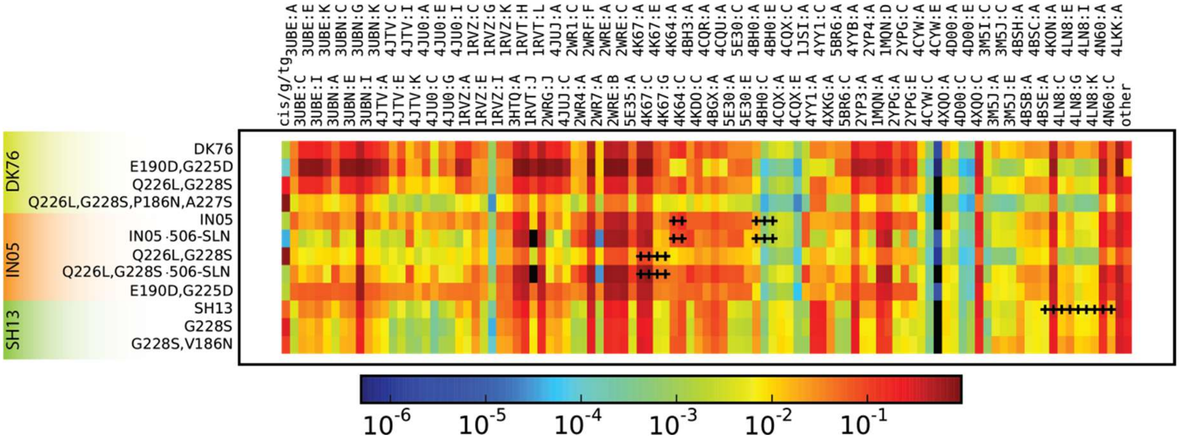


**Figure S4.** The probability that the binding conformation of the receptor analog in the MD simulation is within 1 Å RMSD of the binding conformation in the crystal structures of α2,6-linked receptor analogs. A given human receptor analog bound to any of the simulated HA variants visited almost all the binding conformations observed in crystal structures, even when the crystal structures include HAs of different subtypes, and thus of very different sequences. The plus signs indicate that the crystal structure included HA of the same sequence in the binding pocket as the simulated HA. The first column is the occupancy of the novel conformation, and the last column is the occupancy of other conformations—excluding the novel conformation—that have not yet been observed in other crystallographic structures.

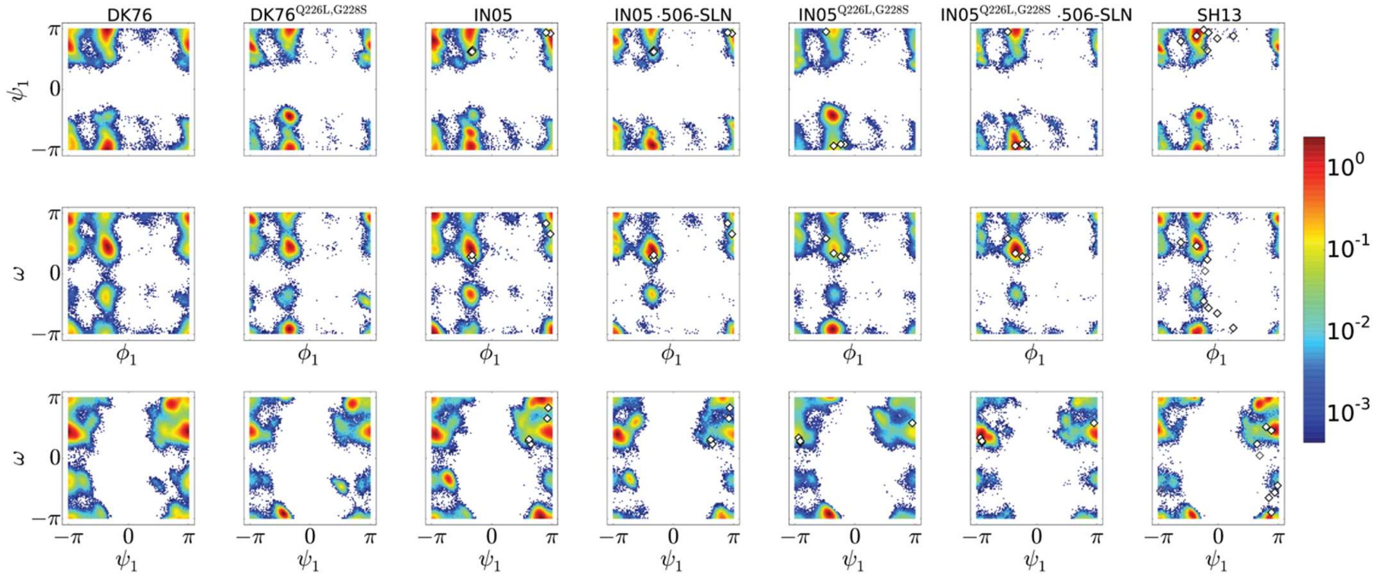


**Figure S5.** The joint distribution in $(\phi_1, \psi_1)$, $(\phi_1, \varpi)$, and $(\psi_1, \varpi)$ for human receptor analogs in complex with a few HA variants. Diamonds indicate the corresponding torsions in the crystal structures of α2,6-linked SAs in complex with HA that have the same binding-pocket amino acid sequence as the simulated HA. The difference in the preferred binding conformations between 6-SLN and 506-SLN in our simulations is supported by the difference seen in the crystal structures of 6-SLN (PDB ID: 4BGX) and 506-SLN (PDB ID: 4K64) in complex with very similar HAs: In 506-SLN bound to the IN05 HA, the glycosidic linkage is in the trans configuration (i.e., $|\varphi_1| > \pi/2$),[2] whereas in 6-SLN bound to the HA of A/Vietnam/1194/2004 (VN04), which differs from the IN05 HA in the binding pocket only at position 193 (it is lysine in VN04 and arginine in IN05; neither makes direct or water-mediated contact with the receptor), the linkage is in the cis configuration (*i.e.* $|\varphi_1| \leq \pi/2$).[3,4] These results, together with the previous crystallographic observation that two different avian receptor analogs bound to H7 HA in two different binding conformations,[5] suggest that the GlcNAc-linked glycan moieties can affect the binding conformations of the three terminal saccharides Neu5Ac, Gal, and GlcNAc.

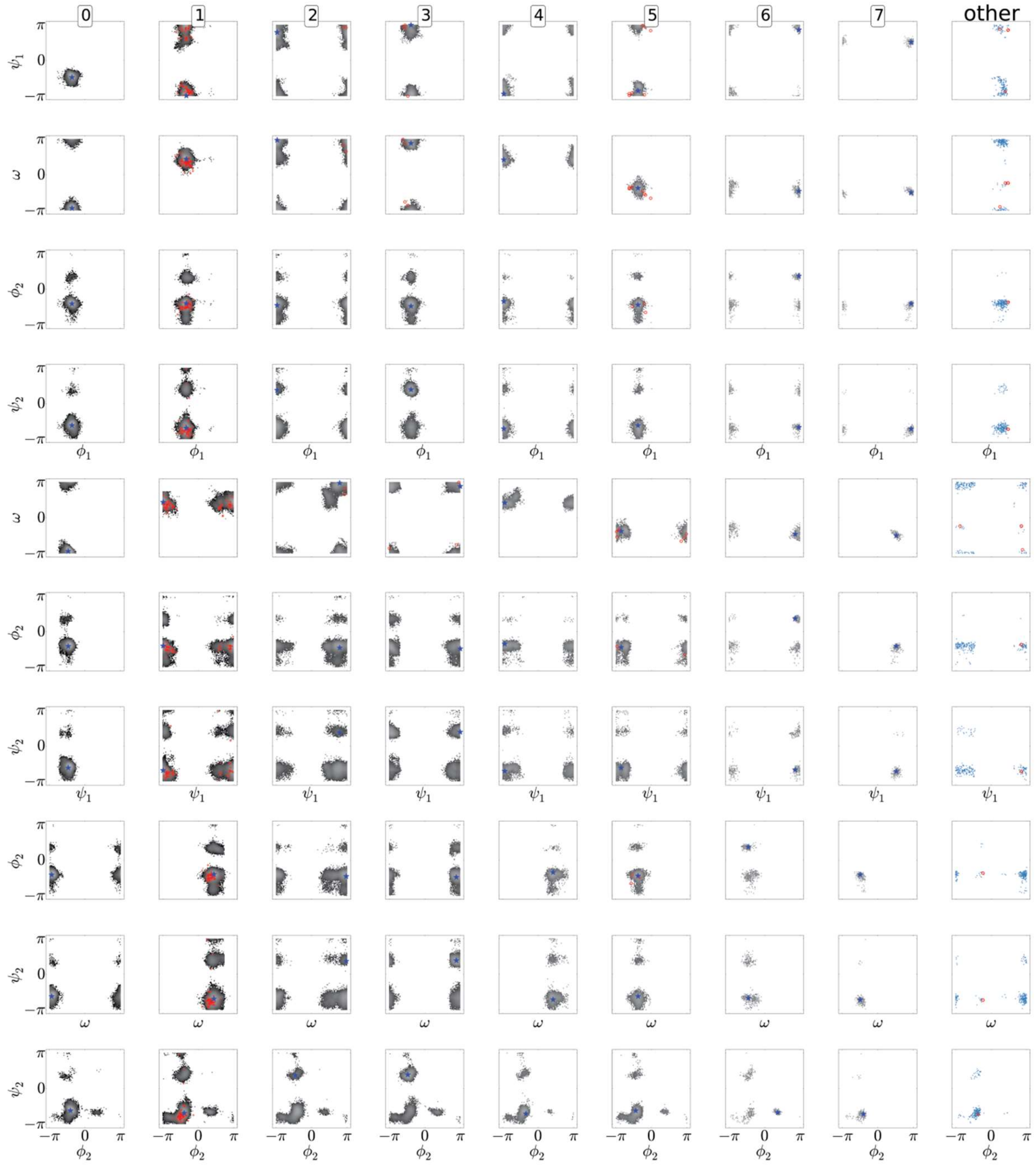


**Figure S6**. The torsional angles sampled by human receptor analogs in different clusters of bound conformations in our simulations. Each row represents a different pair of torsions in $(\phi_1, \psi_1, \varpi, \phi_2, \psi_2)$. The red marks indicate the corresponding torsions in the crystal structures that are assigned to the respective clusters, and the blue diamonds indicate the center of each cluster. Cluster 0 represents the novel conformation, cis/g/tg, shown in Fig. 3b in the main text. "Other" represents transient conformations that are not assigned to any cluster. Three crystal structures are assigned to this cluster of transient conformations: 3M5I (6-SLN bound to HA of the H7N2 strain A/New York/107/2003), one asymmetric unit (I) of 4LN8 (LSTb bound to HA of the H7N9 strain A/Shanghai/2/2013), and 5BR6 (506-SLN bound to HA of the H6N1 strain A/Taiwan/2/2013); they are all sampled, albeit infrequently, in our simulations. Since the clustering was based on similarity in the first three torsions $(\phi_1, \psi_1, \varpi)$, it is clear that each cluster can be further divided into sub-clusters based on similarities in all five torsions. Considering all five torsions resulted in 18 clusters.

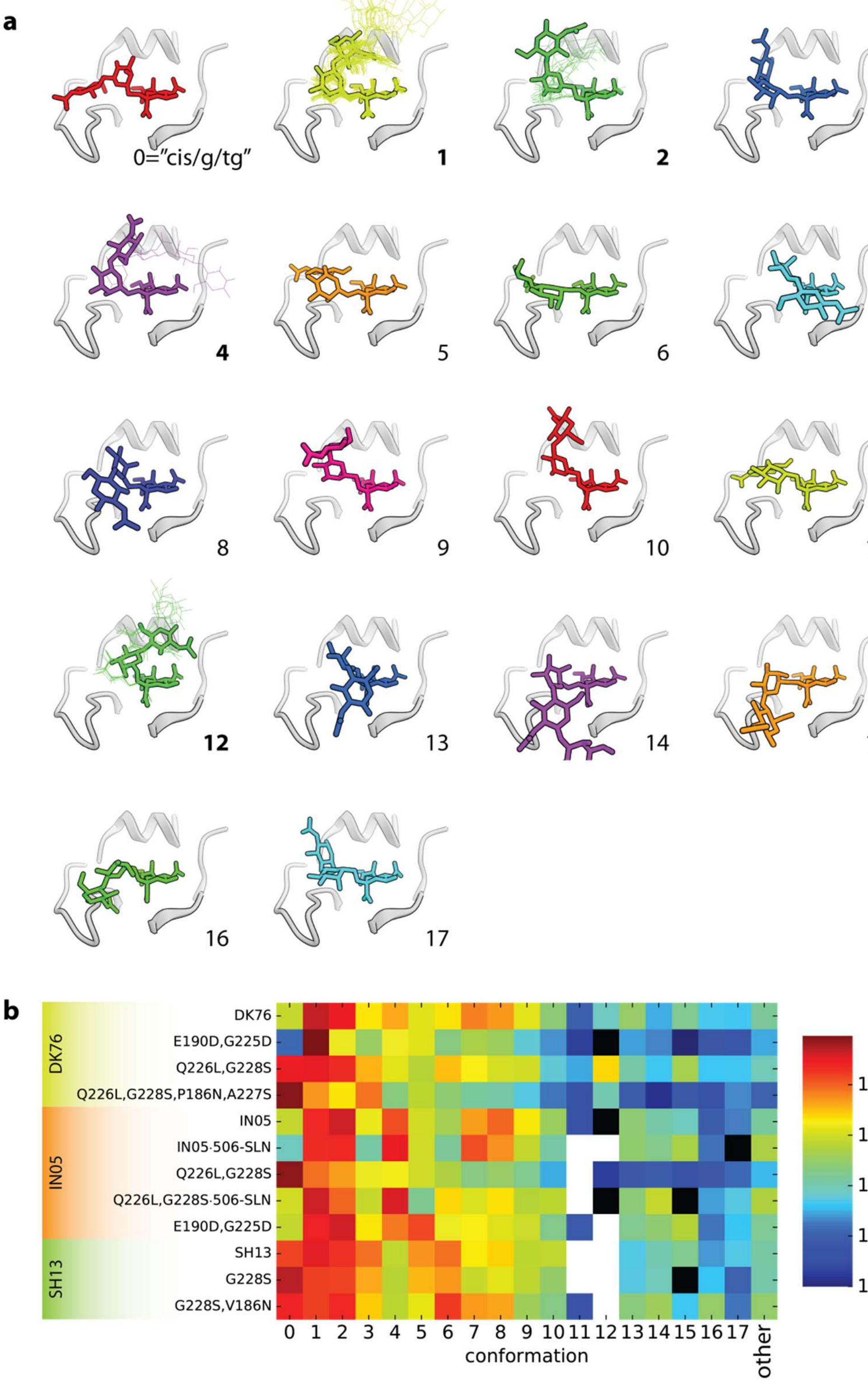


**Figure S7**. The binding conformation of the human receptor analogs 5-SLN and 506-SLN divided into 18 distinct clusters based on the values of the five torsions, $\phi_1, \psi_1, \varpi, \phi_2, \psi_2$, which together determine the positions of Gal and GlcNAc rings. (**a**) Receptor structures corresponding to the centers of the conformation clusters shown in thick lines. Crystal structures in which both Gal and GlcNAc are resolved are shown in thin lines in the corresponding cluster; such crystal structures are represented in only 4 out of 18 clusters (indicated by boldfaced numbers), suggesting that our MD simulations sample more binding conformations than have been determined by crystallography. (**b**) The occupancy of different clusters of conformations by human receptor analogs bound to different HA variants.

**Table S1**. HA variants simulated in this work. HA from three strains, of the H1, H5, and H7 subtypes, are considered: A/duck/Alberta/35/1976 (DK76) of the H1 subtype, A/Indonesia/5/2005 (IN05) of the H5 subtype, and A/Shanghai/2/2013 (SH13) of the H7 subtype.

| **HA variant** | **Amino acid sequence of the molecular construct used in the MD simulations** |
|---|---|
| DK76 | APLQLGKCNVAGWLLGNPECDLLLTANSWSYIIETSNSENGTCYPGEFIDYEELREQLSS |
| | VSSFEKFEIFPKASSWPNHETKGVTAACSYSGASSFYRNLLWITKKGTSYPKLSKSYTNN |
| | KGKEVLVLWGVHHPPSVSEQQSLYQNADAYVSVGSSKYNRRFAPEIAARPKVRGQAGRMN |
| | YYWTLLDQGDTITFEATGNLIAPWYAFALNKGSDSG |
| DK76$^{E190D,G225D}$ | APLQLGKCNVAGWLLGNPECDLLLTANSWSYIIETSNSENGTCYPGEFIDYEELREQLSS |
| | VSSFEKFEIFPKASSWPNHETKGVTAACSYSGASSFYRNLLWITKKGTSYPKLSKSYTNN |
| | KGKEVLVLWGVHHPPSVSDQQSLYQNADAYVSVGSSKYNRRFAPEIAARPKVRDQAGRMN |

YYWTLLDQGDTITFEATGNLIAPWYAFALNKGSDSG

DK76$^{Q226L,G228S}$ APLQLGKCNVAGWLLGNPECDLLLTANSWSYIIETSNSENGTCYPGEFIDYEELREQLSS

VSSFEKFEIFPKASSWPNHETKGVTAACSYSGASSFYRNLLWITKKGTSYPKLSKSYTNN

KGKEVLVLWGVHHPPSVSEQQSLYQNADAYVSVGSSKYNRRFAPEIAARPKVRGLASRMN

YYWTLLDQGDTITFEATGNLIAPWYAFALNKGSDSG

DK76$^{Q226L,G228S,A227S}$ APLQLGKCNVAGWLLGNPECDLLLTANSWSYIIETSNSENGTCYPGEFIDYEELREQLSS

VSSFEKFEIFPKASSWPNHETKGVTAACSYSGASSFYRNLLWITKKGTSYPKLSKSYTNN

KGKEVLVLWGVHHPPSVSEQQSLYQNADAYVSVGSSKYNRRFAPEIAARPKVRGLSSRMN

YYWTLLDQGDTITFEATGNLIAPWYAFALNKGSDSG

DK76$^{Q226L,G228S,P186N,A227S}$ APLQLGKCNVAGWLLGNPECDLLLTANSWSYIIETSNSENGTCYPGEFIDYEELREQLSS

VSSFEKFEIFPKASSWPNHETKGVTAACSYSGASSFYRNLLWITKKGTSYPKLSKSYTNN

KGKEVLVLWGVHHPNSVSEQQSLYQNADAYVSVGSSKYNRRFAPEIAARPKVRGLSSRMN

YYWTLLDQGDTITFEATGNLIAPWYAFALNKGSDSG

IN05

KPLILRDCSVAGWLLGNPMCDEFINVPEWSYIVEKANPTNDLCYPGSFNDYEELKHLLSR

INHFEKIQIIPKSSWSDHEASSGVSSACPYLGSPSFFRNVVWLIKKNSTYPTIKKSYNNT

NQEDLLVLWGIHHPNDAAEQTRLYQNPTTYISIGTSTLNQRLVPKIATRSKVNGQSGRME

FFWTILKPNDAINFESNGNFIAPEYAYKIVKKGDSA

IN05[E190D,G225D]

KPLILRDCSVAGWLLGNPMCDEFINVPEWSYIVEKANPTNDLCYPGSFNDYEELKHLLSR

INHFEKIQIIPKSSWSDHEASSGVSSACPYLGSPSFFRNVVWLIKKNSTYPTIKKSYNNT

NQEDLLVLWGIHHPNDAADQTRLYQNPTTYISIGTSTLNQRLVPKIATRSKVNDQSGRME

| | |
|---|---|
| | FFWTILKPNDAINFESNGNFIAPEYAYKIVKKGDSA |
| IN05$^{Q226L}$ | KPLILRDCSVAGWLLGNPMCDEFINVPEWSYIVEKANPTNDLCYPGSFNDYEELKHLLSR |
| | INHFEKIQIIPKSSWSDHEASSGVSSACPYLGSPSFFRNVVWLIKKNSTYPTIKKSYNNT |
| | NQEDLLVLWGIHHPNDAAEQTRLYQNPTTYISIGTSTLNQRLVPKIATRSKVNGLSGRME |
| | FFWTILKPNDAINFESNGNFIAPEYAYKIVKKGDSA |
| IN05$^{Q226L,G228S}$ | KPLILRDCSVAGWLLGNPMCDEFINVPEWSYIVEKANPTNDLCYPGSFNDYEELKHLLSR |
| | INHFEKIQIIPKSSWSDHEASSGVSSACPYLGSPSFFRNVVWLIKKNSTYPTIKKSYNNT |
| | NQEDLLVLWGIHHPNDAAEQTRLYQNPTTYISIGTSTLNQRLVPKIATRSKVNGLSSRME |
| | FFWTILKPNDAINFESNGNFIAPEYAYKIVKKGDSA |
| SH13 | RTVDLGQCGLLGTITGPPQCDQFLEFSADLIIERREGSDVCYPGKFVNEEALRQILRESG |

GIDKEAMGFTYSGIRTNGATSACRRSGSSFYAEMKWLLSNTDNAAFPQMTKSYKNTRKSP

ALIVWGIHHSVSTAEQTKLYGSGNKLVTVGSSNYQQSFVPSPGARPQVNGLSGRIDFHWL

MLNPNDTVTFSFNGAFIAPDRASFLRGKSMGI

| | |
|---|---|
| SH13$^{G228S,V186N}$ | RTVDLGQCGLLGTITGPPQCDQFLEFSADLIIERREGSDVCYPGKFVNEEALRQILRESG |
| | GIDKEAMGFTYSGIRTNGATSACRRSGSSFYAEMKWLLSNTDNAAFPQMTKSYKNTRKSP |
| | ALIVWGIHHSNSTAEQTKLYGSGNKLVTVGSSNYQQSFVPSPGARPQVNGLSSRIDFHWL |
| | MLNPNDTVTFSFNGAFIAPDRASFLRGKSMGI |
| SH13$^{G228S,V186G}$ | RTVDLGQCGLLGTITGPPQCDQFLEFSADLIIERREGSDVCYPGKFVNEEALRQILRESG |
| | GIDKEAMGFTYSGIRTNGATSACRRSGSSFYAEMKWLLSNTDNAAFPQMTKSYKNTRKSP |
| | ALIVWGIHHSGSTAEQTKLYGSGNKLVTVGSSNYQQSFVPSPGARPQVNGLSSRIDFHWL |

MLNPNDTVTFSFNGAFIAPDRASFLRGKSMGI

| | |
|---|---|
| SH13$^{L226I}$ | RTVDLGQCGLLGTITGPPQCDQFLEFSADLIIERREGSDVCYPGKFVNEEALRQILRESG |
| | GIDKEAMGFTYSGIRTNGATSACRRSGSSFYAEMKWLLSNTDNAAFPQMTKSYKNTRKSP |
| | ALIVWGIHHSVSTAEQTKLYGSGNKLVTVGSSNYQQSFVPSPGARPQVNGISGRIDFHWL |
| | MLNPNDTVTFSFNGAFIAPDRASFLRGKSMGI |

**Table S2.** The population of cis/trans configurations of the first glycosidic bond. The glycosidic bond is considered to be in the cis-configuration if $|\varphi_1| \leq \pi/2$.

| | cis fraction in MD | configuration in crystal structure |
|---|---|---|
| 6-SLN in solution | 0.88 ± 0.01 | |
| 506-SLN in solution | 0.863 ± 0.005 | |
| 503-SLN in solution | 0.760 ± 0.007 | |
| DK76 + 6-SLN | 0.622 ± 0.004 | unresolved (2WRH) |
| $DK76^{E190D,G225D}$ + 6-SLN | 0.976 ± 0.001 | |
| $DK76^{Q226L,G228S}$ + 6-SLN | 0.70 ± 0.02 | |
| $DK76^{Q226L,G228S,P186N,A227S}$ + 6-SLN | 0.972 ± 0.002 | |
| IN05 + 6-SLN | 0.49 ± 0.01 | cis (4BH0; tyTy has same binding site sequence as IN05) |
| IN05 + 506-SLN | 0.61 ± 0.04 | trans (4K64) |
| VN04 + 6-SLN | 0.42 ± 0.02 | cis (4BGX) |

| | | |
|---|---|---|
| $IN05^{Q226L,G228S}$ + 6-SLN | 0.94 ± 0.01 | |
| $IN05^{Q226L,G228S}$ + 506-SLN | 0.87 ± 0.02 | cis (4K67) |

**Table S3**. HA variants characterized for receptor analog binding by MST assay in this work.

| **HA variant** | **Amino acid sequence of the molecular construct used in the MST measurements** |
| --- | --- |
| DK76 | MEAKLFVLFCTFTVLKADTICVGYHANNSTDTVDTVLEKNVTVTHSVNLLEDS<br>HNGKLCSLNGIAPLQLGKCNVAGWLLGNPECDLLLTANSWSYIIETSNSENGT<br>CYPGEFIDYEELREQLSSISSFEKFEIFPKASSWPNHETTKGVTAACSYSGAS<br>SFYRNLLWITKKGTSYPKLSKSYTNNKGKEVLVLWGVHHPPSVSEQQSLYQNA<br>DAYVSVGSSKYNRRFAPEIAARPKVRGQAGRMNYYWTLLDQGDTITFEATGNL<br>IAPWYAFALNKGSDSGIITSDAPVHNCDTRCQTPHGALNSSLPFQNVHPITIG<br>ECPKYVKSTKLRMATGLRNVPSIQSRGLFGAIAGFIEGGWTGMIDGWYGYHHQ<br>NEQGSGYAADQKSTQNAIDGITNKVNSVIEKMNTQFTAVGKEFNNLERRIENL<br>NKKVDDGFLDVWTYNAELLVLLENERTLDFHDSNVRNLYEKVKSQLRNNAKEI<br>GNGCFEFYHKCDDECMESVKNGTYDYPKYSEESKLNREEIDGVKLESMGVYQI<br>LAIYSTVASSLVLLVSLGAISFWMCSNGSLQCRICI |
| DK76$^{E190D,G225D}$ | MEAKLFVLFCTFTVLKADTICVGYHANNSTDTVDTVLEKNVTVTHSVNLLEDS<br>HNGKLCSLNGIAPLQLGKCNVAGWLLGNPECDLLLTANSWSYIIETSNSENGT<br>CYPGEFIDYEELREQLSSISSFEKFEIFPKASSWPNHETTKGVTAACSYSGAS<br>SFYRNLLWITKKGTSYPKLSKSYTNNKGKEVLVLWGVHHPPSVSDQQSLYQNA<br>DAYVSVGSSKYNRRFAPEIAARPKVRDQAGRMNYYWTLLDQGDTITFEATGNL<br>IAPWYAFALNKGSDSGIITSDAPVHNCDTRCQTPHGALNSSLPFQNVHPITIG<br>ECPKYVKSTKLRMATGLRNVPSIQSRGLFGAIAGFIEGGWTGMIDGWYGYHHQ<br>NEQGSGYAADQKSTQNAIDGITNKVNSVIEKMNTQFTAVGKEFNNLERRIENL<br>NKKVDDGFLDVWTYNAELLVLLENERTLDFHDSNVRNLYEKVKSQLRNNAKEI<br>GNGCFEFYHKCDDECMESVKNGTYDYPKYSEESKLNREEIDGVKLESMGVYQI<br>LAIYSTVASSLVLLVSLGAISFWMCSNGSLQCRICI |

DK76$^{Q226L,G228S}$ MEAKLFVLFCTFTVLKADTICVGYHANNSTDTVDTVLEKNVTVTHSVNLLEDS
HNGKLCSLNGIAPLQLGKCNVAGWLLGNPECDLLLTANSWSYIIETSNSENGT
CYPGEFIDYEELREQLSSISSFEKFEIFPKASSWPNHETTKGVTAACSYSGAS
SFYRNLLWITKKGTSYPKLSKSYTNNKGKEVLVLWGVHHPPSVSEQQSLYQNA
DAYVSVGSSKYNRRFAPEIAARPKVRGLASRMNYYWTLLDQGDTITFEATGNL
IAPWYAFALNKGSDSGIITSDAPVHNCDTRCQTPHGALNSSLPFQNVHPITIG
ECPKYVKSTKLRMATGLRNVPSIQSRGLFGAIAGFIEGGWTGMIDGWYGYHHQ
NEQGSGYAADQKSTQNAIDGITNKVNSVIEKMNTQFTAVGKEFNNLERRIENL
NKKVDDGFLDVWTYNAELLVLLENERTLDFHDSNVRNLYEKVKSQLRNNAKEI
GNGCFEFYHKCDDECMESVKNGTYDYPKYSEESKLNREEIDGVKLESMGVYQI
LAIYSTVASSLVLLVSLGAISFWMCSNGSLQCRICI

DK76$^{Q226L,G228S,A227S}$ MEAKLFVLFCTFTVLKADTICVGYHANNSTDTVDTVLEKNVTVTHSVNLLEDS
HNGKLCSLNGIAPLQLGKCNVAGWLLGNPECDLLLTANSWSYIIETSNSENGT
CYPGEFIDYEELREQLSSISSFEKFEIFPKASSWPNHETTKGVTAACSYSGAS
SFYRNLLWITKKGTSYPKLSKSYTNNKGKEVLVLWGVHHPPSVSEQQSLYQNA
DAYVSVGSSKYNRRFAPEIAARPKVRGLSSRMNYYWTLLDQGDTITFEATGNL
IAPWYAFALNKGSDSGIITSDAPVHNCDTRCQTPHGALNSSLPFQNVHPITIG
ECPKYVKSTKLRMATGLRNVPSIQSRGLFGAIAGFIEGGWTGMIDGWYGYHHQ
NEQGSGYAADQKSTQNAIDGITNKVNSVIEKMNTQFTAVGKEFNNLERRIENL
NKKVDDGFLDVWTYNAELLVLLENERTLDFHDSNVRNLYEKVKSQLRNNAKEI
GNGCFEFYHKCDDECMESVKNGTYDYPKYSEESKLNREEIDGVKLESMGVYQI
LAIYSTVASSLVLLVSLGAISFWMCSNGSLQCRICI

DK76$^{Q226L,G228S,P186N}$ MEAKLFVLFCTFTVLKADTICVGYHANNSTDTVDTVLEKNVTVTHSVNLLEDS
HNGKLCSLNGIAPLQLGKCNVAGWLLGNPECDLLLTANSWSYIIETSNSENGT
CYPGEFIDYEELREQLSSISSFEKFEIFPKASSWPNHETTKGVTAACSYSGAS

SFYRNLLWITKKGTSYPKLSKSYTNNKGKEVLVLWGVHHPNSVSEQQSLYQNA
DAYVSVGSSKYNRRFAPEIAARPKVRGLASRMNYYWTLLDQGDTITFEATGNL
IAPWYAFALNKGSDSGIITSDAPVHNCDTRCQTPHGALNSSLPFQNVHPITIG
ECPKYVKSTKLRMATGLRNVPSIQSRGLFGAIAGFIEGGWTGMIDGWYGYHHQ
NEQGSGYAADQKSTQNAIDGITNKVNSVIEKMNTQFTAVGKEFNNLERRIENL
NKKVDDGFLDVWTYNAELLVLLENERTLDFHDSNVRNLYEKVKSQLRNNAKEI
GNGCFEFYHKCDDECMESVKNGTYDYPKYSEESKLNREEIDGVKLESMGVYQI
LAIYSTVASSLVLLVSLGAISFWMCSNGSLQCRICI

DK76$^{Q226L,G228S,P186N,A227S}$

MEAKLFVLFCTFTVLKADTICVGYHANNSTDTVDTVLEKNVTVTHSVNLLEDS
HNGKLCSLNGIAPLQLGKCNVAGWLLGNPECDLLLTANSWSYIIETSNSENGT
CYPGEFIDYEELREQLSSISSFEKFEIFPKASSWPNHETTKGVTAACSYSGAS
SFYRNLLWITKKGTSYPKLSKSYTNNKGKEVLVLWGVHHPNSVSEQQSLYQNA
DAYVSVGSSKYNRRFAPEIAARPKVRGLSSRMNYYWTLLDQGDTITFEATGNL
IAPWYAFALNKGSDSGIITSDAPVHNCDTRCQTPHGALNSSLPFQNVHPITIG
ECPKYVKSTKLRMATGLRNVPSIQSRGLFGAIAGFIEGGWTGMIDGWYGYHHQ
NEQGSGYAADQKSTQNAIDGITNKVNSVIEKMNTQFTAVGKEFNNLERRIENL
NKKVDDGFLDVWTYNAELLVLLENERTLDFHDSNVRNLYEKVKSQLRNNAKEI
GNGCFEFYHKCDDECMESVKNGTYDYPKYSEESKLNREEIDGVKLESMGVYQI
LAIYSTVASSLVLLVSLGAISFWMCSNGSLQCRICI

IN05

MEKIVLLLAIVSLVKSDQICIGYHANNSTEQVDTIMEKNVTVTHAQDILEKTH
NGKLCDLDGVKPLILRDCSVAGWLLGNPMCDEFINVPEWSYIVEKANPTNDLC
YPGSFNDYEELKHLLSRINHFEKIQIIPKSSWSDHEASSGVSSACPYLGSPSF
FRNVVWLIKKNSTYPTIKKSYNNTNQEDLLVLWGIHHPNDAAEQTRLYQNPTT
YISIGTSTLNQRLVPKIATRSKVNGQSGRMEFFWTILKPNDAINFESNGNFIA
PEYAYKIVKKGDSAIMKSELEYGNCNTKCQTPMGAINSSMPFHNIHPLTIGEC
PKYVKSNRLVLATGLRNSPQRESRRKKRGLFGAIAGFIEGGWQGMVDGWYGYH

HSNEQGSGYAADKESTQKAIDGVTNKVNSIIDKMNTQFEAVGREFNNLERRIE
NLNKKMEDGFLDVWTYNAELLVLMENERTLDFHDSNVKNLYDKVRLQLRDNAK
ELGNGCFEFYHKCDNECMESIRNGTYNYPQYSEEARLKREEISGVKLESIGTY
QILSIYSTVASSLALAIMMAGLSLWMCSNGSLQCRICI

IN05$^{Q226L,G228S}$ MEKIVLLLAIVSLVKSDQICIGYHANNSTEQVDTIMEKNVTVTHAQDILEKTH
NGKLCDLDGVKPLILRDCSVAGWLLGNPMCDEFINVPEWSYIVEKANPTNDLC
YPGSFNDYEELKHLLSRINHFEKIQIIPKSSWSDHEASSGVSSACPYLGSPSF
FRNVVWLIKKNSTYPTIKKSYNNTNQEDLLVLWGIHHPNDAAEQTRLYQNPTT
YISIGTSTLNQRLVPKIATRSKVNGLSSRMEFFWTILKPNDAINFESNGNFIA
PEYAYKIVKKGDSAIMKSELEYGNCNTKCQTPMGAINSSMPFHNIHPLTIGEC
PKYVKSNRLVLATGLRNSPQRESRRKKRGLFGAIAGFIEGGWQGMVDGWYGYH
HSNEQGSGYAADKESTQKAIDGVTNKVNSIIDKMNTQFEAVGREFNNLERRIE
NLNKKMEDGFLDVWTYNAELLVLMENERTLDFHDSNVKNLYDKVRLQLRDNAK
ELGNGCFEFYHKCDNECMESIRNGTYNYPQYSEEARLKREEISGVKLESIGTY
QILSIYSTVASSLALAIMMAGLSLWMCSNGSLQCRICI

IN05$^{Q226L,G228S,S227A}$ MEKIVLLLAIVSLVKSDQICIGYHANNSTEQVDTIMEKNVTVTHAQDILEKTH
NGKLCDLDGVKPLILRDCSVAGWLLGNPMCDEFINVPEWSYIVEKANPTNDLC
YPGSFNDYEELKHLLSRINHFEKIQIIPKSSWSDHEASSGVSSACPYLGSPSF
FRNVVWLIKKNSTYPTIKKSYNNTNQEDLLVLWGIHHPNDAAEQTRLYQNPTT
YISIGTSTLNQRLVPKIATRSKVNGLASRMEFFWTILKPNDAINFESNGNFIA
PEYAYKIVKKGDSAIMKSELEYGNCNTKCQTPMGAINSSMPFHNIHPLTIGEC
PKYVKSNRLVLATGLRNSPQRESRRKKRGLFGAIAGFIEGGWQGMVDGWYGYH
HSNEQGSGYAADKESTQKAIDGVTNKVNSIIDKMNTQFEAVGREFNNLERRIE
NLNKKMEDGFLDVWTYNAELLVLMENERTLDFHDSNVKNLYDKVRLQLRDNAK
ELGNGCFEFYHKCDNECMESIRNGTYNYPQYSEEARLKREEISGVKLESIGTY
QILSIYSTVASSLALAIMMAGLSLWMCSNGSLQCRICI

**Table S4.** Comparison of observables of solution conformations between MD calculations and NMR measurements for trisaccharides. 6-SLN was simulated in the MD calculations, but sialyl-α(2,6)-lactose was used in NMR experiments.[6] 6-SLN and sialyl-α(2,6)-lactose differ only in the last sugar at the reducing end: It is a acetylglucosamine in 6-SLN, and a glucose in sialyl-α(2,6)-lactose. The results from MD simulations are in good agreement with NMR measurements: the root-mean-square error (RMSE) in the NOE distances is 0.48 Å, and RMSE in the $^3J$ coupling constants, which are computed from parameterized Karplus equations,[6–9] is 0.7 Hz.

| *NOE distances (Å)* | | *NMR* | $\langle r^{-6}\rangle_{\mathrm{MD}}^{-1/6}$ |
|---|---|---|---|
| **Neu5Ac / Gal** | $H3_{ax}$ / H6R | 3.3 | 3.44 ± 0.03 |
| | $H3_{ax}$ / H6S | 3.3 | 3.27 ± 0.03 |
| | $H3_{ax}$ / H4 | >5 | 4.26 ± 0.01 |
| | $H3_{eq}$ / H6R | 5 | 4.236 ± 0.003 |
| | $H3_{eq}$ / H6S | 4 | 4.25 ± 0.02 |
| | $H3_{eq}$ / H5 | 5 | 4.812 ± 0.007 |
| | $H3_{eq}$ / H4 | >5 | 4.30 ± 0.03 |
| | OH8 / H6R | 3.5 | 5.12 ± 0.04 |
| | OH8 / H6S | >4 | 4.51 ± 0.06 |

| | | | |
|---|---|---|---|
| | OH7 / Glc O3 | 3 | 3.27 ± 0.04 |
| | C2 / H6R | 2.6 | 2.749 ± 0.004 |
| | C2 / H6S | >2.6 | 2.764 ± 0.003 |
| **Gal / Gal** | H4 / H6R | 3.3 | 2.91 ± 0.01 |
| | H4 / H6S | 2.6 | 3.51 ± 0.02 |
| | **Gal / Glc** H1 / H4 | 2.5 | 2.368 ± 0.004 |
| | H1 / H6S | 3.5 | 3.17 ± 0.03 |
| | H6R / OH3 | 3.5 | 3.850 ± 0.009 |
| | H6R / O3 | - | 4.43 ± 0.01 |
| | H6S / O3 | - | 4.06 ± 0.01 |
| *coupling constants $^3J$ (Hz)* | | *NMR* | *MD* |
| **Neu5Ac / Gal** | C2 / H6S | 1.8 (2) | 2.16 ± 0.03 |
| | C2 / H6R | 2.8 (2) | 2.72 ± 0.02 |

| | | | |
|---|---|---|---|
| **Gal / Gal** | H5 / H6R | 8.5 (2) | 10.1 ± 0.2 |
| | H5 / H6S | 3.7 (2) | 4.0 ± 0.1 |
| | C4 / H6R | 2.9 (5) | 3.24 ± 0.01 |
| | C4 / H6S | 1.8 (5) | 1.90 ± 0.04 |
| | **Gal / Glc** H1 / C4 | 4.2 (2) | 2.937 ± 0.006 |
| | C1 / H4 | 5.1 (4) | 5.26 ± 0.01 |

**Table S5.** Comparison of observables of solution conformations between MD calculations and NMR measurements[10] for 506-SLN (LSTc) at 295 K. The RMSE in the NOE distances is 0.3 Å, and the RMSE in the $^3J$ coupling constants is 0.4 Hz.

| *NOE distances (Å)* | *NMR* | $\langle r^{-6} \rangle_{\mathrm{MD}}^{-1/6}$ |
|---|---|---|
| **Neu5Ac / Gal1** H5 / GlcNAc $CH_3$ | 4.24 | 4.58 ± 0.04 |
| **GlcNAc / Gal2** H1 / H3 | 1.94 | 2.316 ± 0.002 |
| H1 / H4 | 3.05 | 3.162 ± 0.007 |
| **Gal2 / Glc** H1 / H4 | 1.96 | 2.366 ± 0.002 |
| H1 / H5 | | |

| *coupling constants $^3J$ (Hz)* | | *NMR* | *MD* |
|---|---|---|---|
| **Neu5Ac / Gal1** | C2 / H6R | 2.6 | 2.73 ± 0.02 |
| **Gal1 / Gal1** | H5 / H6R | 10 | 9.88 ± 0.09 |
| **Gal1 / GlcNAc** | H1 / C4 | 3.6 | 2.99 ± 0.01 |

| | | |
|---|---|---|
| **GlcNAc / Gal2** H1 / C3 | 4.1 | 3.387 ± 0.005 |
| C1 / H3 | 4.4 | 4.188 ± 0.009 |
| **Gal2 / Glc** H1 / C4 | 3.4 | 2.95 ± 0.02 |

**Table S6.** Comparison of observables of solution conformations between MD calculations and NMR measurements[10] for 503-SLN (LSTa) at 295K. The RMSE in the NOE distances is 0.5 Å, and the RMSE in $^3J$ coupling constants is 0.7 Hz. The only substantial disagreement between the MD and the NMR results lies in the relative configurations of Gal2 and Glc, the last two sugars at the reducing end. This disagreement suggests that the force field we used may not accurately describe the interactions across the glycosidic bond between Gal2 and Glc. Since we simulated the HA binding of the trisaccharide 3-SLN, which does not contain these two moieties, our results and conclusion are unlikely affected by such force field inaccuracies.

| *NOE distances (Å)* | | *NMR* | $\langle r^{-6} \rangle_{\mathrm{MD}}^{-1/6}$ |
|---|---|---|---|
| **Neu5Ac / Gal1** | $H3_{ax}$ / H3 | 2.76 | 3.05 ± 0.02 |
| | $H3_{eq}$ / H3 | 4.70 | 4.22 ± 0.01 |
| | H5 / GlcNAc $CH_3$ | - | 6.6 ± 0.1 |
| **Gal1 / GlcNAc** | H1 / H3 | 2.25 | 2.422 ± 0.009 |
| | H1 / $CH_3$ | 4.00 | 3.43 ± 0.04 |
| **GlcNAc / Gal2** | H1 / H3 | 1.95 | 2.341 ± 0.004 |
| | H1 / H4 | 2.84 | 3.146 ± 0.004 |
| **Gal2 / Glc** | | 2.20 | 2.365 ± 0.003 |

| | | |
|---|---|---|
| H1 / H4 | | |
| H1 / H5 | 2.80 | 3.92 ± 0.03 |
| *coupling constants ³J (Hz)* | *NMR* | *MD* |
| **Neu5Ac / Gal1** C2 / H3 | 4.4 | 3.96 ± 0.04 |
| **Gal1 / GlcNAc** H1 / C3 | 2.6 | 2.94 ± 0.02 |
| C1 / H3 | 5.4 | 4.61 ± 0.09 |
| **GlcNAc / Gal2** H1 / C3 | 4.2 | 3.39 ± 0.02 |
| C1 / H3 | 4.5 | 4.18 ± 0.02 |
| **Gal2 / Glc** H1 / C4 | 4.1 | 2.96 ± 0.02 |

**Movie S1**. MD simulation of the human receptor analog, 6-SLN, binding to the SH13$^{L226I}$ HA, which has the same amino acid sequence as the HA of the A/Hangzhou/1/2013 strain. In the 4.13-μs simulation, the receptor analog, shown in sticks, spontaneously bound to the HA three times, and unbound from the HA twice. The Neu5Ac moiety is shown in orange. The crystal structure of 6-SLN in complex with A/Netherlands/219/2003 HA (PDB ID: 4DJ8), of very similar sequence, is briefly displayed in gray to show the agreement between the complex structure generated by the MD simulation and the crystal structure. The intermolecular hydrogen bonds that are observed in all crystal structures of HA in complex with receptors are subsequently highlighted by purple lines, with key residues of the HA shown in ball-and-stick representation. On binding to HA the second time, the receptor first bound with Neu5Ac outside the pocket, but changed to the crystallographic pose after ~1 μs. When the receptor was bound to HA, it interconverted among a wide range of conformations, and its contacts with HA changed dynamically, as shown in the last part of the movie, with HA represented by a gray molecular surface, with the atoms forming instantaneous contacts highlighted in yellow.
